# Targeting Myc activates a tissue-specific tumour resolution programme

**DOI:** 10.64898/2026.02.06.704498

**Authors:** Roderik M. Kortlever, Tania Campos, Stefan Boeing, Kazimir Uzwyshyn-Jones, Alessandra Perfetto, Gerard I. Evan

## Abstract

Neoplastic transformation parallels hallmark cellular programs of tissue regeneration and wound repair^1,2^. However, the mechanisms driving tumour regression upon oncogenic driver inhibition—and why it often fails—remain poorly understood^3–7^. Oncogenic KRas mutations and Myc deregulation, two archetypal cancer drivers, frequently occur and cooperate to promote aggressive lung adenocarcinoma (LUAD)^8–12^. To investigate the mechanistic consequences of targeting Myc in LUAD, we applied spatiotemporally controlled genetic and functional perturbations in a reversible KRas/Myc-driven mouse model, integrated with RNA sequencing and immune protein profiling of tumours and their microenvironment. Acute oncogenic Myc inactivation in epithelial tumour cells elicits a localised regenerative immune response crucially dependent on rapid, transient release of the alarmin cytokine interleukin-33 (IL-33) by alveolar type 2 tumour cells. As a sentinel signal for Myc loss, IL-33 signalling reverses tumour immunosuppression and neoangiogenesis, critically recruits eosinophils, and promotes neoplastic cell elimination, driving regression beyond mere growth arrest. Notably, brief systemic recombinant IL-33 administration to mice with KRas/Myc-driven LUAD induces robust eosinophil influx and near-complete tumour resolution. Together, these findings demonstrate that blocking Myc activates an innate, tissue-intrinsic immune programme rooted in resolution of wound repair and capable of driving regression when activated in a tumour. This opens the possibility of treating cancer not only by blocking mitogenic oncogenic drivers but also by pro-actively triggering pro-resolution pathways.

## Introduction

Targeted cancer therapies, despite often eliciting an initial response, frequently fail to produce durable therapeutic outcomes in the main due to extensive inherent and adaptive redundancy in many signalling pathways^3,4,13^. We also lack a comprehensive understanding of how and why blockading oncogenic signals, which are naturally turning on and off all the time to regulate cell proliferation in normal tissues, elicits tumour cell death, a still largely unexplained phenomenon dubbed oncogene addiction. We clearly need targets that have minimal biological redundancy, that are conduits that lie downstream of the kinase and G protein “oncogenome”, and whose blockade leads to cancer cell death via a well-documented mechanism. Aside its current state as undruggable, Myc fits the bill perfectly. The oncogenic capacity of oncogenic KRas and Myc, either each alone or in synergy, serve as paradigmatic models for elucidating oncogene-driven tumour cell growth and death, across a wide range of mouse models of cancer^5,14–18^. Their deregulation via mutations or relentless activation by upstream growth factor signalling directly programmes multiple neoplastic growth promoting processes, including cellular proliferation, metabolism, angiogenesis, inflammation, and the immune microenvironment^19–22^. Of note, we have previously demonstrated that the tumorigenic programmes driven by the same KRas/Myc combination drive very distinct oncogenic phenotypes in different tissues. Thus, in mouse models of cancer progression, combination of the identical duo of KRas and Myc activation induces lung adenocarcinoma (LUAD) in the lung and pancreatic ductal adenocarcinoma (PDAC) in the pancreas^12,21^: both lesion types recapitulating the signature histopathological and molecular features of their spontaneous human counterparts. Conversely, acute Myc withdrawal in established LUAD or PDAC triggers tumour regression and, to an approximation, re-establishment of original tissue architecture. The therapeutic potential and any clinical efficacy of inhibiting either oncogene is widely anticipated^6,7^. Ras and Myc are well-recognised as critical drivers and essential mediators of tumour progression and maintenance in mouse models of lung cancer^10,11,23–25^. When cooperatively activated in distal airway epithelial cells they generate a complex and structured lesion identical to that of spontaneous lung adenocarcinoma - angiogenic, high microvessel density, AT2-cell lepidic growth, immune infiltration, invasion of alveoli^12^. Reversibly, selective loss of deregulated Myc in the tumour epithelial cells of our LUAD mouse model instantly triggers profound microenvironmental changes leading to tumour regression^12^. This process bears striking similarities to tissue regeneration following injury: regression and resolution of neoplastic lesions share molecular components, pathways, and biological mechanisms with wound healing^26–30^, a concept long recognised^1,31^. Prompted by the rapidity of the regression pathology and analogy to damage resolution in normal tissue, we hypothesised that targeting a driver oncogene may actively programme tumour regression by activating pathways involved in resolving tissue damage, restoring the lung towards its homeostatic state.

## Results

### Acute loss of oncogenic Myc in LUAD triggers instant tumour cell IL-33 production

We previously used a mouse model of non-small cell lung adenocarcinoma (LUAD) driven by a KRas*^G12D^* allele together with a reversibly switchable 4-OHT-dependent allele of the Myc oncoprotein (MycER^T2^) - *LSL-KRas^G12D^; ROSA26-Lox-STOP-Lox-MycER^T^*^2^ (*KM* mouse) - to assess the role of sustained Myc to maintain LUAD. This showed that acute downregulation of Myc (hereafter referred to as “Myc-OFF”) triggers rapid and synchronous onset of regression of established LUAD across the multiplicity of independent lung tumour foci generated in each individual mouse^12^ . Importantly, even though Myc is de-activated only in the epithelial tumour cells, what follows is rapid recruitment of a complex choreography of many microenvironmental cell types that, in turn, drives highly organized interactions between inflammatory, immune, vascular and mesenchymal cells. Within a few hours of Myc-OFF, tumour cells drop out of cycle and start to die, angiogenesis shuts down, CD3+ T and NKp46+ NK cells (both of which are rapidly expelled from incipient KRas*^G12D^*-driven lung tumours when Myc is initially switched on) rapidly re-infiltrate the tumours, while CD206^+^ macrophages are ejected from them^12^. The immediacy of onset, synchrony and interwoven complexity of Myc-OFF-driven LUAD regression all suggest that regression is not merely passive reversal of tumourigenesis but a dedicated, tissue morphogenic programme. The regression programme is initiated by a fall in intracellular Myc activity, specifically within the epithelial LUAD cells, and then propagated extracellularly, presumably through the release of instructive signal cascades that, in turn, commission the many cell types that run the regression programme. To investigate the early transcriptional and cell dynamics that are immediately engaged in LUAD after switching Myc-OFF, we first established the rapidity with which Myc transcriptional activity decays in the LUAD tumours upon Myc withdrawal. *KM* mice were transferred to a tamoxifen diet for 4 weeks, long enough to establish significant MycER^T2^- driven multifocal lung adenocarcinoma growth, along with all its signature phenotypic features (inflammation, angiogenesis, local invasion, immune suppression, etc.)^12^. We then transferred the *KM* mice back to normal food to switch MycER^T2^ OFF for various times. To monitor how rapidly Myc transcriptional activity attenuates after Myc-OFF, we used bulk RNA-sequencing at 0-, 2- and 4-hours post-tamoxifen withdrawal (Extended Data Fig. 1a) to follow the expression of documented Myc gene targets in *KM* lung tissue (Human Molecular Signatures Database (MSigDB) Hallmark collection M5926 (^32^)). By 2 hours post-transfer to tamoxifen- free food, and further pronounced by 4 hours, expression of almost all Myc target genes had significantly fallen (Extended Data Fig. 2a, b and Extended Data Table 1). By contrast, a selection of “housekeeping” genes known to not be direct Myc targets^33^ showed no overall directional change in their expression in the first few hours following Myc-OFF (Extended Data Fig. 2c and Extended Data Table 1). To gain some overall sense of the impact of Myc-OFF on transcriptional programmes classically associated with key cellular pathways we analysed changes in gene sets from the MSigDB Hallmark collection. Four hours after switching Myc-OFF (4 hrs Myc-OFF vs Myc-ON), most pathway gene sets that were significantly enriched for when Myc was ON for 4 weeks (Myc-ON vs KRas-only) had significantly reversed their expression (Extended Data Fig. 2d). Thus, attenuation of Myc transcriptional activity follows very rapidly after deactivation of MycER^T2^ in tumour cells and this is accompanied by downregulation of mRNA in Myc target genes and Myc activity-associated cellular pathways. To confirm the kinetics and consequence of Myc-OFF in lung, we isolated RNA from tumour-laden *KM* mouse lungs in which Myc had been continuously active for 4 weeks (‘0’ hours Myc-OFF), along with equivalent lungs collected at 2, 4, 8, 12, and 24 hours after Myc-OFF. These were then subjected to single cell RNA sequence analysis (Extended Data Fig.1b). Across all cell types and time points, we identified some 30 distinct cell types (Extended Data Fig. 2e, f, Extended Data Table 2) of which only four epithelial cell types showed expression of MycER^T2^ – mucinous ciliated epithelial cells, Alveolar Type 1 (AT1), cuboidal Alveolar Type 2 (AT2), and Club cells. We then used gene set enrichment analysis (GSEA) to confirm rapid downregulation of Myc target genes (evident within 2 hours of Myc-OFF) in these four lung epithelial cell types (Extended Data Fig. 2g) following withdrawal of tamoxifen.

### IL-33 is rapidly and transiently released in response to Myc OFF

In our *KM* LUAD mice, Myc is toggled on and off only in sporadic lung epithelial tumour cells. Nonetheless, Myc-OFF has very rapid and widespread impacts on many other cell types within the lung tumours. Based on this observation, we reasoned that in response to Myc-OFF tumour cells must engage a downstream signal pathway cascade that instructs the regression process.

To identify the earliest instructive signals engaged after switching Myc-OFF in LUAD, we first used tissue immune arrays to probe protein extracts derived from tumour-laden lungs at various times after Myc-OFF – 12hrs, 1, 2, 3 and 7 days (Extended Data Fig. 1c, Extended Data Fig. 3a). By 12 hours post Myc-OFF, the cytokine IL-33 was the most significantly elevated signalling molecule (Fig. 1a) although this was transient and had subsided by 3 days post Myc-OFF (Extended Data Fig. 3b). We next used differential gene expression (DGE) using a collection of 350 immune-signalling genes to interrogate signalling genes expressed in the MycER^T2^-positive epithelial cell types during a 24-hour Myc OFF time course (Extended Data Table 3). The Alveolar Type 2 (AT2) cell population, alone, uniquely showed expression and enrichment for the RNA encoding the alarmin interleukin 33 (IL-33) (Fig. 1b, Extended Data Fig. 3c). Immunohistochemistry confirmed expression of the IL-33 in the nuclei of tumour AT2 epithelial cells (Fig. 1c, d). IL-33 has established credentials as an alarmin sentinel that coordinates regeneration in lung and other tissues after tissue injury^34–36^, we therefore addressed whether IL-33 is causally required for tumour regression.

**Figure 1.**
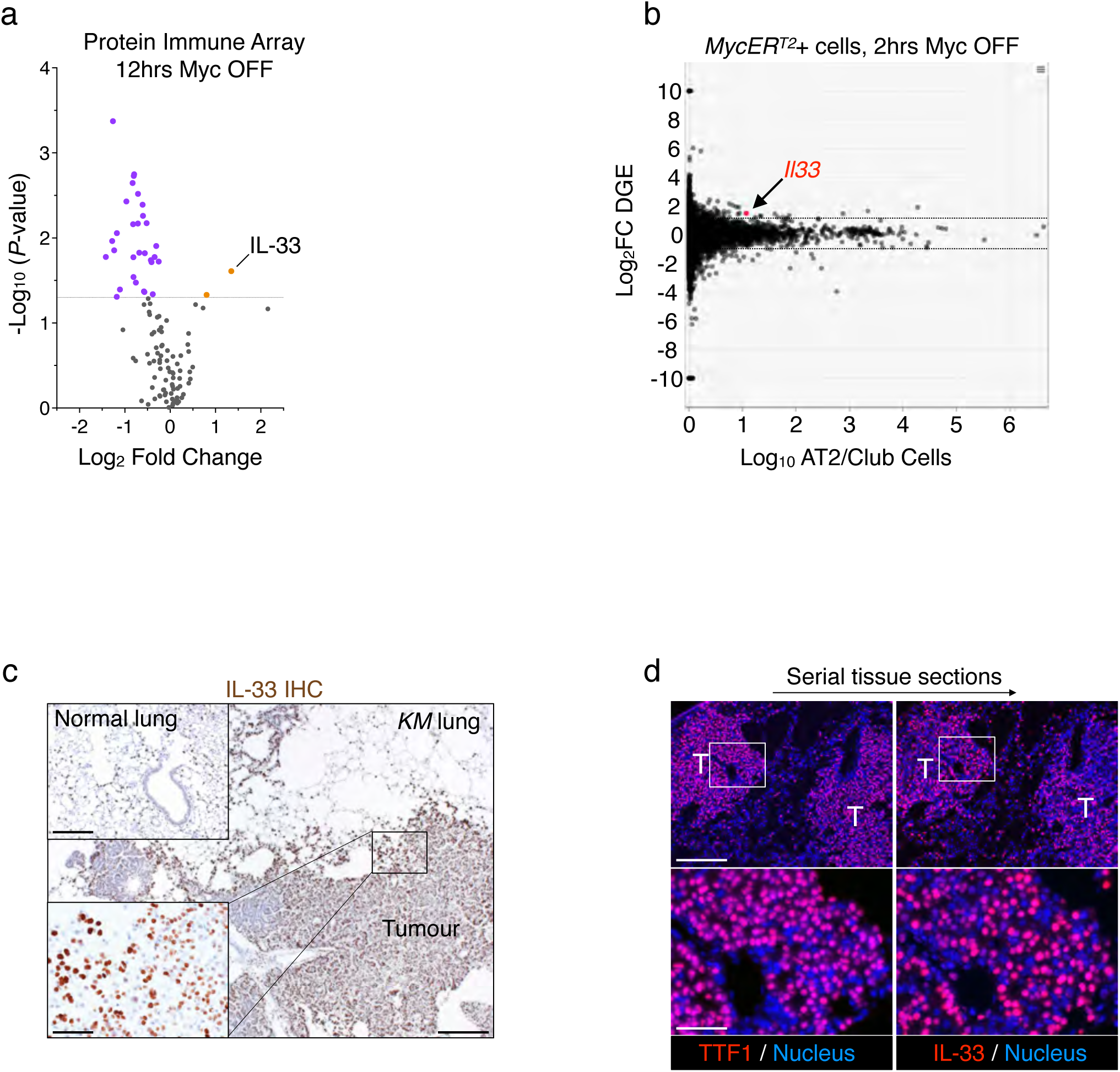
Acute loss of deregulated Myc initiates immediate epithelial tumour cell IL-33 production. **a**, Induction of IL33 protein levels after Myc-OFF. The Volcano plot depicts fold protein expression 12 hours after Myc OFF (compared with Myc ON), as analysed by immune cytokine array. Significantly up- and downregulated proteins are shown in tangerine and grape colour, respectively. Each individual datapoint represents the median value for a specific protein. Data are from three independent biological replicate experiments. Statistical analysis performed using multiple unpaired *t*-tests with Welch’s correction on Myc OFF 12 hrs (n = 8) versus Myc ON (n = 9) comparison. **b**, Induction of *Il33* transcription levels in MycER^T2^-positive AT2 cells after Myc-OFF. Differential gene expression (DGE) of the *MycER^T^*^2^-positive fraction of the AT2/Club cell cluster in tumour-laden lungs 2 hours post-tamoxifen (2hrs Myc OFF vs Myc ON). Data are generated from 4-8 mouse per condition from up to 4 independent biological replicate experiments. **c**, IL33 is expressed in LUAD tumour cells. Representative qualitative immunohistochemistry (IHC) of protein IL-33 in the lung and tumours of *KM* mice after no tumour induction (‘normal lung’ inset top left) versus one week of tamoxifen treatment (Myc ON, ‘*KM* lung’). Scale bars: IHC sections normal and *KM* lung 200μm, *KM* lung inset 50μm. **d**, IL33 is expressed in tumour AT2 epithelial tumour cells. Representative qualitative immunofluorescence of protein IL-33 and AT2 cell type marker TTF1 expression in the tumours of *KM* mice after two weeks of tamoxifen treatment (Myc ON). Bottom row is an enlarged insets from top row. T = tumour. Scale bars: sections top row 200μm, bottom row 50μm.

### IL-33 signalling is required for Myc OFF induced tumour regression

To determine directly whether IL-33 release is causally required for Myc-OFF-driven tumour regression, we repeated the previous Myc OFF time course while blocking IL-33 signalling systemically throughout the first week post Myc-OFF with blocking antibodies specific for the non-redundant IL-33 receptor ST2. Control animals were treated with sub-type matched IgG (Extended Data Fig.1d). As previously reported^12^, in IgG treated control *KM* mice Myc-OFF triggered rapid onset of macroscopic tumour regression, greatly reducing tumour burden by 7 days. In sharp contrast, no observable reduction in overall tumour burden was evident in Myc-OFF tumour-bearing *KM* animal treated with anti-ST2 blocking antibody (Fig. 2a). Hence, IL- 33 signalling plays a necessary immediate early role in LUAD to engage the downstream Myc-OFF-dependent regression. As outlined above, Myc-OFF in the LUAD tumour cell compartment triggers a complex morphogenic and dynamic regression programme characterised by onset of tumour cell cycle arrest and death, rapid tumoral influx of CD3+T cells and NKp46+ NK cells, efflux of CD206^+^ macrophages, and shutdown of angiogenesis^12^. Of these attributes, IL-33-ST2 blockade profoundly inhibited Myc-OFF-induced tumour cell death, influx of CD3+ T and NK cell into tumours, and shutdown of angiogenesis (Fig. 2b-d, Extended Data Fig. 4a). By contrast, Myc-OFF abrogated tumour cell proliferation irrespective of whether IL-33 signalling was blocked (Fig. 2b), consistent with the well-documented cell-intrinsic role of Myc in driving cell proliferation directly. Myc-OFF-induced efflux of CD206+ macrophages from tumours was likewise unaffected by IL-33-ST2 blockade (Fig. 2b). B220+ B cell influx into tumours after Myc-OFF was modestly impeded by IL-33-ST2 blockade (Extended Data Fig. 4b). Of the granulocyte lineage, Myc-OFF induced influx of EPX+ eosinophil influx into the tumours was prevented by IL-33-ST2 blockade while Ly6B.2+ neutrophil efflux was unperturbed (Extended Data Fig. 4b). Overall, we conclude that inhibition of IL-33 signalling selectively impedes some, but not all, aspects of the tumour regression programme triggered by Myc downregulation.

**Figure 2.**
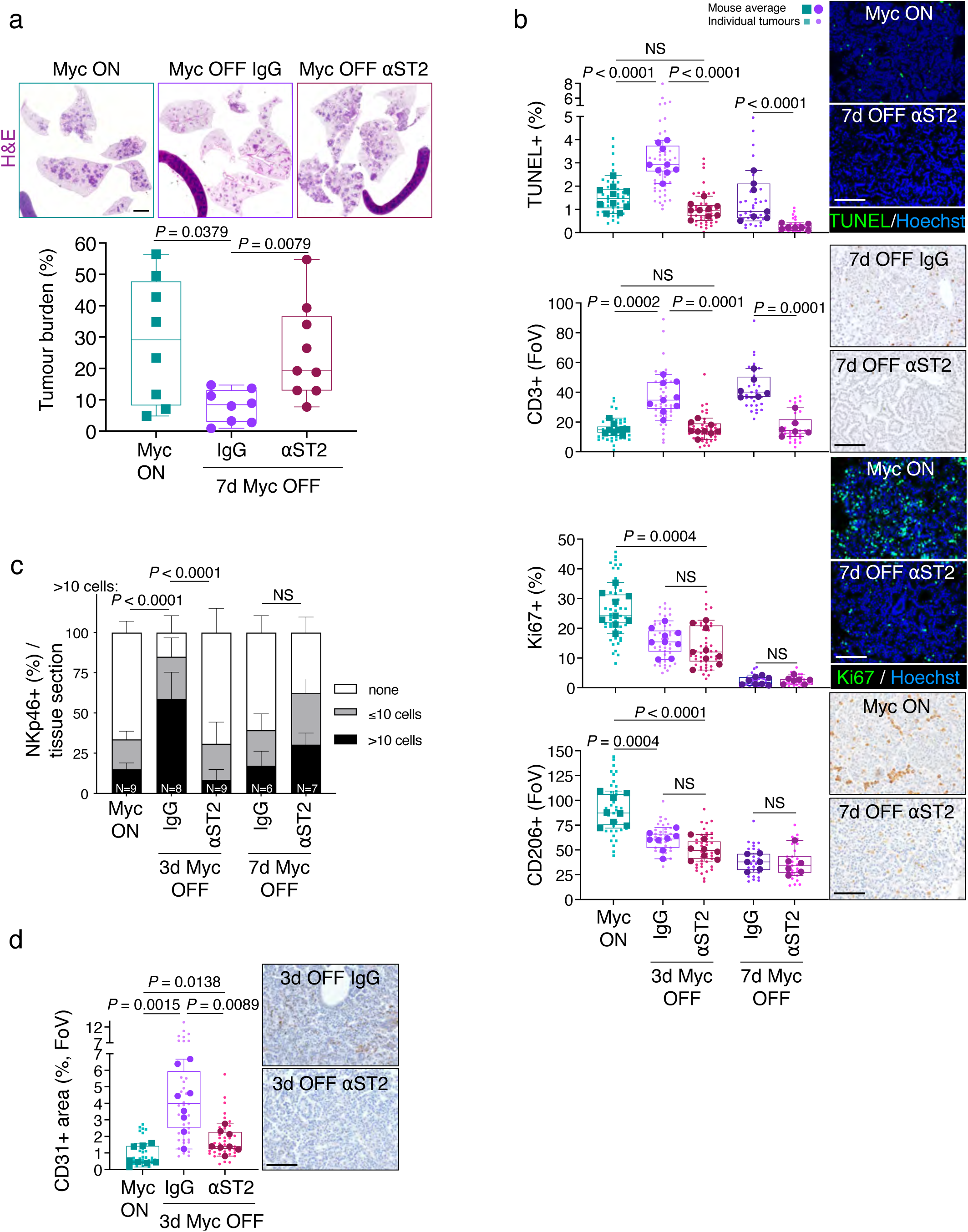
IL-33 Signalling is necessary for tumour regression and regulates immune-suppression and vascular normalisation after Myc OFF. **a**, Blocking IL33 receptor ST2 after Myc-OFF prevents tumour regression. Quantification of tumour burden in *KM* mice treated with tamoxifen (Myc ON), followed by cessation of treatment for one week while simultaneously treated with either control IgG or blocking antibody against IL-33 receptor ST2. Associated, above: representative H&E stain of whole lung sections for either condition / treatment. Scale bar: 2mm. n = 8 or 9 mice per condition. Statistical analysis was performed using the multiple Mann-Whitney tests. Data are from 3 independent biological replicate experiments. **b**, IL33 signalling after Myc-OFF is required for LUAD tumour cell death and local immunity. Quantification of immunohistochemical analysis for cell death (TUNEL), T cells (CD3), proliferation (Ki67), and Macrophages (CD206) in lung tumours in *KM* mice treated with tamoxifen (Myc ON) followed by Myc OFF for 3 or 7 days while simultaneously treated with anti-ST2 or IgG control antibody. FoV = field of view. Panels to their right: representative immunohistochemistry. n = 6 to 9 mice per condition or timepoint. Statistical analysis was performed using the multiple unpaired *t*-tests with Welch’s correction and based on the mouse averages. Scale bars 100μm. **c**, Blocking IL33 receptor ST2 after Myc-OFF prevents NK cell attraction. Quantification of immunohistochemical analysis for tumour-juxtaposed NK cells (NKp46) of lung tumours in *KM* mice treated with tamoxifen (Myc ON), followed by Myc OFF for 3 or 7 days while simultaneously treated with blocking anti-ST2 versus IgG control antibody. n = 6 to 9 mice per condition or timepoint. Statistical analysis was performed using the two-way analysis of variance (ANOVA) with Tukey’s multiple comparisons test. **d**, Blocking IL33 receptor ST2 after Myc-OFF prevents normalisation of neoangiogenesis. Quantification of immunohistochemical analysis for vascular marker CD31 in *KM* mice treated with tamoxifen (Myc ON) followed by Myc OFF for 3 days while simultaneously treated with anti-ST2 or IgG control antibody. FoV = field of view. n = 8 mice per condition or timepoint. Statistical analysis was performed using the multiple unpaired *t*-tests with Welch’s correction and based on the mouse averages. Scale bar 100μm. **b-d**, Data are from 2 (3 days Myc OFF) or 3 (7 days Myc OFF) independent biological replicate experiments.

### AT2 epithelial lung tumour cells are the source of IL-33 driving lung tumour regression

Like most signalling molecules, IL-33 signalling is highly contextual and dependent on cell source, signal intensity and timing, duration, and modulation by concurrent meta-signals. However, if IL-33 release is the instructive signal triggered by Myc-OFF, we would expect it to originate in the LUAD tumour cells themselves. To confirm that the source of the initiating IL-33 downstream regression signal downstream of Myc-OFF originates in the MycER^T2^ positive AT2 tumour cells (see Fig. 1d), we crossed our *KM* mice into the *Il33^fl/fl^-eGFP* background. In *Il33^fl/fl^-eGFP* mice, *loxP* sites flank exons 5-7 of the endogenous *interleukin 33* (*Il33*) gene. In addition, an EGFP “hit-and-run” CRE reporter lies in the 3’ UTR. In these *R26-LSL-MER;LSL-KrasG12D;Il33-Flox-eGFP* (*KMI*) mice, AdV-CRE focal activation of Cre recombinase in lung epithelium not only co-activates expression of both KRas*^G12D^* and MycER^T2^ but simultaneously deletes expression of functional IL-33, while marking the recombined epithelial cell and its progeny with EGFP. *KMI* mice were then treated with tamoxifen for two weeks to activate MycER^T2^ and drive formation of multiple LUAD. At this time point (i.e. prior to Myc-OFF) there were no measurable differences in overall tumour burdens in mice with one versus two floxed *Il33* alleles (Fig. 3a), indicating that IL-33 expression is not necessary for KRas*^G12D^* and Myc-driven tumour growth and maintenance. Tamoxifen was then withdrawn (Myc-OFF) for a week (Extended data Fig.1e). Tumours in control mice in which conditional IL-33 deletion was hemizygous (*KMI^fl/+^*) underwent rapid regression. By contrast, tumours from *KMI^fl/fl^* mice exhibit no regression one week after Myc-OFF (Fig. 3a). We established that in *KMI^fl/fl^* mice ∼90% of the tumours were IL-33 negative (Fig. 3b, c; Extended Data Fig. 5a). We next ascertained in more detail cell cycle, cell death and immune cell changes in the first 3 and 7 days after switching Myc OFF in *KMI^fl/+^* versus *KMI^fl/fl^* mice. In accordance with blocking IL-33 receptor ST2 *in KM* mice, after switching Myc OFF the absence of tumour cell IL-33 in *KMI^fl/fl^* mice did not impair the downregulation of tumour proliferation nor macrophage efflux, yet did prevent tumour cell death and CD3+ T cell, NK cell, and eosinophil influx (Fig. 3d, Extended Data Fig. 5b, c; see also Fig. 2b, c and Extended Data Fig 4b). We conclude that tumour regression following Myc-OFF is completely dependent on the IL-33 specifically released from the MycER^T2^ positive tumour epithelial cells.

**Figure 3.**
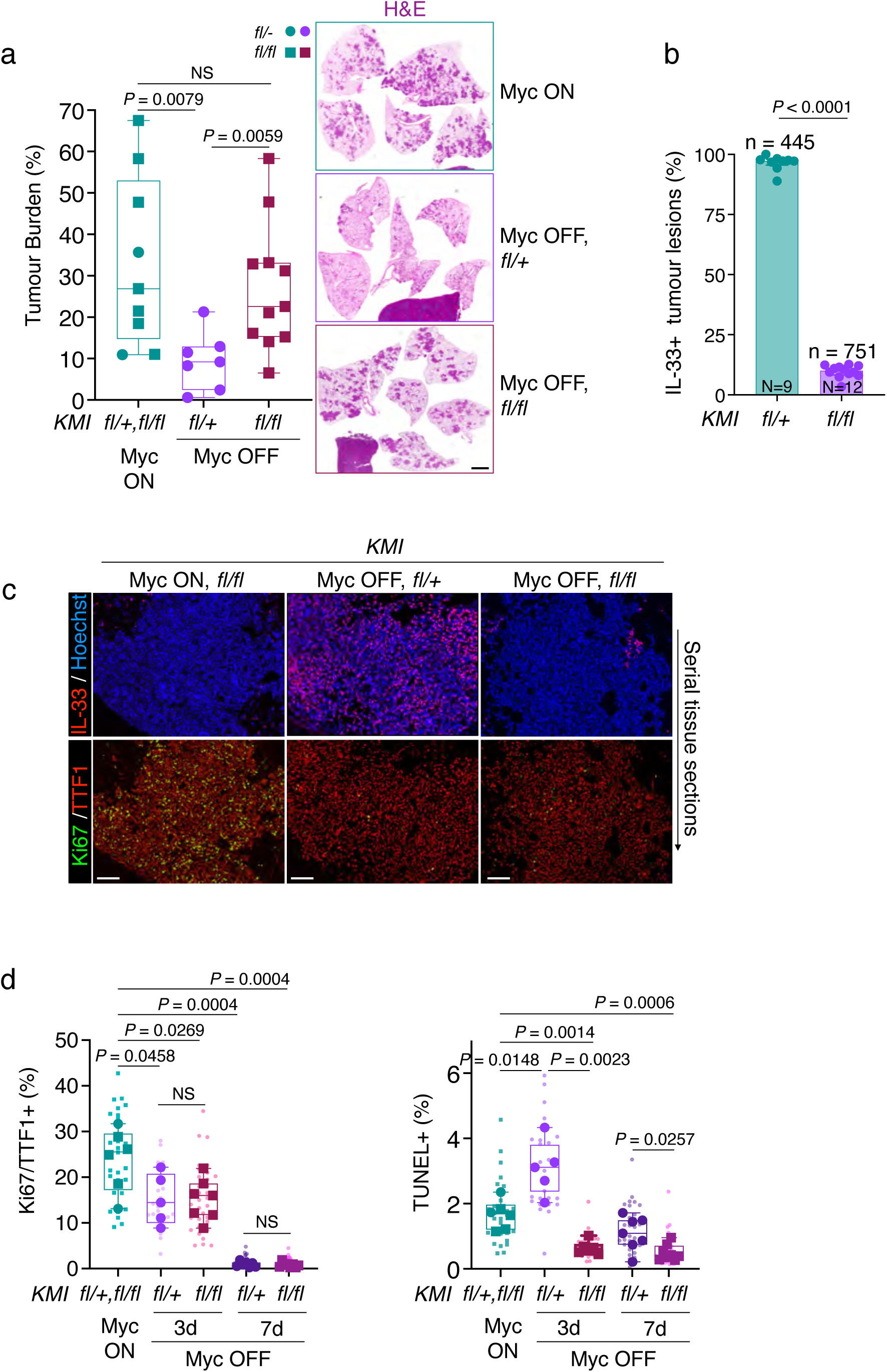
AT2 epithelial lung tumour cells are the IL-33 source for tumour regression. **a**, Absence of IL33 in AT2 tumour cells prevents regression after Myc-OFF. Quantification of tumour burden in hemizygous (*fl/+*, circular datapoints) and homozygous conditional IL-33 null (*fl/fl*, square datapoints) *KMI* littermate mice treated with tamoxifen (Myc ON), followed by cessation of treatment for one week (no tamoxifen; Myc OFF) in either *KMI^fl/+^* or *KMI^fl/fl^*mice. Associated, to the right: representative H&E stain of whole lung sections for either genotype / treatment. Scale bar: 2mm. n = 7-11 mice per condition. Statistical analysis was performed using the multiple Mann-Whitney tests. Data are from 2 independent biological replicate experiments. **b**, Efficiency of selective loss of *Il33* in *KMI* mice. Quantification of tumour lesion IL-33 positivity as measured by immunofluorescence in either *KMI^fl/+^* or *KMI^fl/fl^*mice. N = number of mice assessed and n = number of tumours assessed per genotype; either *fl/+* (N = 9 mice; n = 445 tumours) or *fl/fl* (N = 12 mice, n = 751 tumours), for both Myc ON and Myc OFF conditions. Data are from 2 independent biological replicate experiments. **c**, *KMI^fl/fl^* mice express no measurable IL33 in their AT2 tumour cells. Representative immunofluorescence analysis of protein IL-33 (top row) and lung epithelial cell type AT2 marker protein TTF1 combined with proliferation marker Ki67 (bottom row) in tumours of *KMI* mice at indicated conditions (Myc ON or OFF) and genotypes (*fl/+* or *fl/fl)*. Scale bars 100μm. **d**, Absence of tumour cell death after Myc-OFF in *KMI^fl/fl^* mice. Quantification of immunohistochemical analysis for proliferating AT2 cells (Ki67/TTF1) or cell death (TUNEL) of lung tumours in *KMI^fl/+^* or *KMI^fl/fl^*mice treated with tamoxifen (Myc ON) followed by regular food (Myc OFF) for 3 or 7 days. n = 5 to 9 mice per condition and timepoint. Data are from 2 independent biological replicate experiments. **b**, **d**: Statistical analysis was performed using the multiple unpaired *t*-tests with Welch’s correction and based on the mouse averages.

### An ectopic systemic pulse of IL-33 is sufficient to drive near-complete tumour regression

Given that IL-33 is rapidly and transiently produced by LUAD cells upon Myc-OFF (see Fig. 1a, Extended Data Fig. 3b) and that this is required for LUAD tumour regression (see Fig. 2a, 3a), we asked whether an externally administered pulse of IL-33 in KRas*^G12D^*/MycER^T2^-driven adenocarcinomas would suffice to trigger drive tumour regression. To investigate this, MycER^T2^ was activated in *KM* mice for two weeks to establish lungs laden with rapidly progressing adenocarcinomas. Mice were then given two intraperitoneal doses of recombinant IL-33 (rIL-33) spaced 24 hrs apart and then maintained on tamoxifen food for a further 2 weeks to maintain oncogenic Myc activity (Extended Data Fig. 1f). This rIL-33 regimen was chosen to mimic both the 24-48 hr peak of IL-33 expression following acute lung damage or tissue trauma^37–39^ and to mirror the *in vivo* timing and peak levels of IL-33 induced by Myc-OFF (see Extended Data Fig. 3b). Compared with mice in which Myc-ON activity was maintained for 2 or 4 weeks, the tumour burden in the rIL-33 treated mice was profoundly reduced and, indeed, even lower than that of the KRas*^G12D^*-only driven adenoma burden (Fig. 4a). In rIL-33 treated mice we also observed occasional damaged alveolar regions, intrabronchiolar epithelial hyperplasia and pockets of eosinophils in lungs of rIL-33 treated mice post two weeks rIL-33 treatment (Fig. 4a; see arrows of the H&E insets). This prompted us to investigate the status of rIL-33-treated mice and their tumours in more detail when changing in the first 3 and 7 days after rIL-33 treatment (Extended Data Fig.1g). In the first week after rIL-33 treatment tumour burden also fell very immediately and very rapidly after rIL-33 treatment (Extended Data Fig. 6a) and this was associated with a marked increase in tumour cell death together with decrease of proliferation (Fig 4b, c). We also noted a marked increase in inter- and intra-tumoral presence of eosinophils (Fig 4d, e). To address whether this post rIL-33 treatment eosinophil influx was a Myc-ON or LUAD-associated phenotype, or a generic IL-33-triggered type 2 immunity response in the lung^40^ we treated *KM* mice with and without tumours with rIL-33 and determined the levels of eosinophils in either condition (Extended Data Fig. 1h). Enhanced eosinophil influx was prominent at 7 days post rIL-33 administration and independent of tumour presence: systemic rIL-33 treatment of *KM* mice devoid of lung tumours also provoked a strong eosinophil presence (Extended Data Fig. 6b). We conclude that transient recombinant IL-33 treatment is sufficient to impose resolution of LUAD cells accompanied by an elevated high presence of eosinophils.

**Figure 4.**
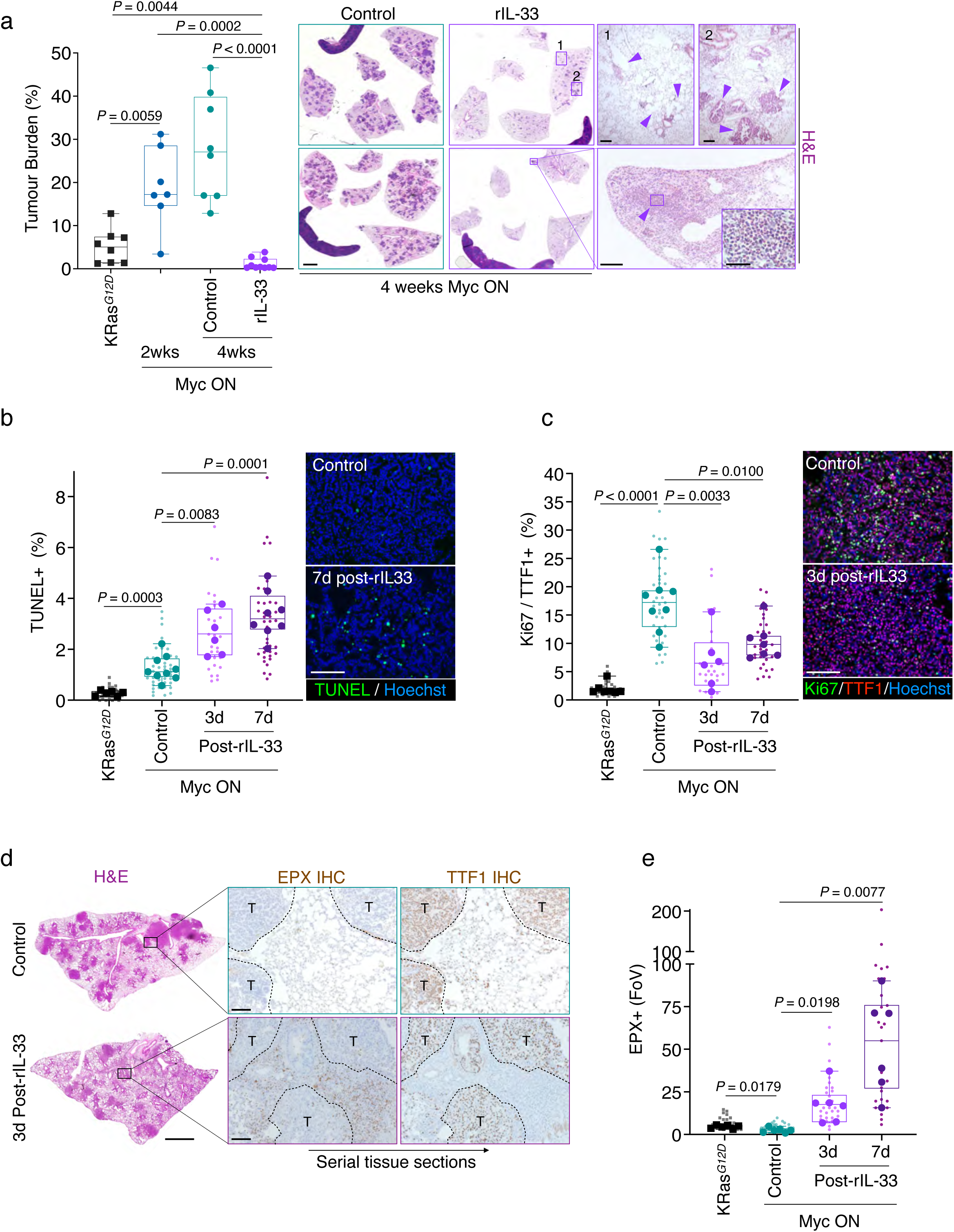
Transient IL-33 treatment is sufficient to impose LUAD resolution. **a**, Exogenous IL33 imposes tumour resolution. Quantification of tumour burden in *KM* mice without treatment (KRas*^G12D^-*only), treated with tamoxifen (Myc ON) for 2 weeks, or Myc ON for 4 weeks total with either PBS (control) or recombinant IL33 (rIL-33) treatment after 2 weeks. Associated, to the right: representative H&E staining of whole lung sections plus enlargements (insets 1 and 2 top row, to the right), or one small part of one lung lobe (inset bottom row, to the right) to highlight remnant lesions (see arrows) or eosinophil aggregation (see inset), respectively. n = 7 to 10 mice per condition/timepoint. Scale bars: whole lung sections 2mm.; top right panels 1, 2: 200μm; bottom right panel 100μm, inset 20μm. Statistical analysis was performed using the multiple Mann-Whitney tests. **b**, rIL33 induces tumour cell death. Quantification of immunohistochemical analysis for cell death (TUNEL) of lung tumours in *KM* mice non-treated (KRas*^G12D^*-only), treated with tamoxifen (Myc ON), or 3- or 7-days post systemic rIL-33 treatment while Myc ON. Panels to the right: representative immunohistochemistry. n = 6 to 9 mice per condition/timepoint. Scale bar 100μm. **c**, rIL33 reduces tumour cell proliferation. Quantification of immunohistochemical analysis for proliferation in AT2 cells (Ki67/TTF1) of lung tumours in non-treated *KM* mice (KRas*^G12D^*- only) versus *KM* mice treated with tamoxifen (Myc ON), or 3- or 7-days post systemic rIL-33 treatment while Myc ON. Panels to the right: representative immunohistochemistry. n = 6 to 9 mice per condition/timepoint. Scale bar 100μm. **d**, rIL33 induces a dramatic eosinophil influx in the lung. Representative H&E stains of a large lung lobe tissue section 3 days post-control or post-rIL-33 treatment (see Extended Data Fig. 1g and 4a, b). Scale bar 2mm. Insets enhanced to the right: representative immunohistochemistry (IHC) staining for eosinophils (EPX) or AT2 cells (TTF1) of serial tissue sections. T = tumour. Scale bars 100μm. **e**, rIL33 enriches for intratumoral eosinophils. Quantification of immunohistochemical analysis of eosinophil presence (EPX) in lung tumours of *KM* mice non-treated (KRas*^G12D^-* only), treated with tamoxifen (Myc ON, control), or 3- or 7-days post systemic rIL-33 treatment while continued Myc ON. FoV = field of view. n = 6 to 9 mice per condition/timepoint. **b**, **c**, **e**: Statistical analysis was performed using the multiple unpaired *t*-tests with Welch’s correction and based on the mouse averages. **a, b, c, e**: Data are from 3 independent biological replicate experiments.

### Eosinophils are required for Myc OFF driven, and IL-33 mediated, tumour regression

This prompted us to investigate whether eosinophils too are necessary for tumour regression downstream of Myc-OFF. First, we confirmed expression of the IL-33 specific receptor ST2 on eosinophils^41^ (Fig. 5a). To directly address whether eosinophils are functionally involved in Myc OFF regression downstream of IL-33, we crossed our *KM* mice into mice with a targeted deletion of the high-affinity GATA binding site in the GATA1 promoter (ΔdblGATA) that is crucial for the development of eosinophils^42,43^, resulting in *LSL-KRas^G12D^; ROSA26-Lox-STOP-Lox-MycER^T^*^2^; Δ*dblGATA* (*KMG)* mice. We then subjected both heterozygous and homozygous null littermates to Myc-ON/OFF treatment to compare their tumour growth and regression dynamics with those of *KM* mice (Extended Data Fig. 1i). Myc-induced tumour progression was similar in *KM* versus *KMG^null^* mice: by comparison, Myc-OFF-induced tumour regression was completely dependent on eosinophils and absent in *KMG^null^* mice, as compared to *KM* or *KMG^het^* mice after Myc-OFF (Fig. 5b). As expected, absence of eosinophils did not abrogate the profound suppression of proliferation observed in Myc-OFF tumours, but profoundly did profoundly suppress tumour cell death Myc-OFF-induced tumour cell death (Fig. 5c). Furthermore, the influx of CD3+ T and NK cells that Myc-OFF normally induces was entirely blocked in *KMG^null^* mice but the efflux of CD206^+^ macrophages was unaffected, just as was seen with IL-33-dependence manner in *KM* and *KMI^fl/fl^* mice. (Extended Data Fig. 7a, b; see Fig. 2b, c, and Extended Data Fig 5b, c). We conclude that eosinophils play an obligate part of the regression programme downstream of Myc-OFF.

**Figure 5.**
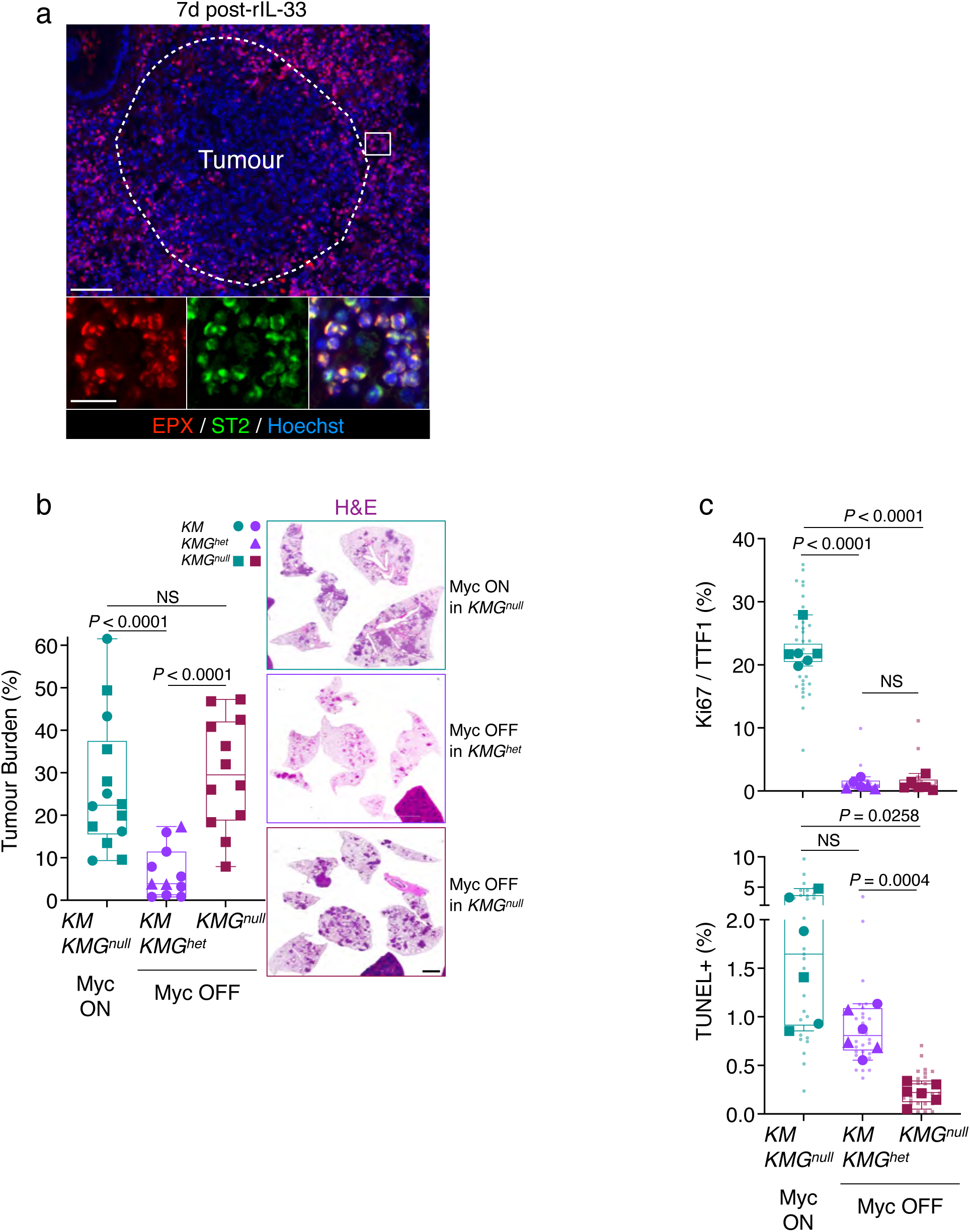
Eosinophils are necessary for Myc OFF driven tumour regression. **a**, Eosinophils express IL33-receptor ST2. Representative qualitative immunofluorescence analysis of cells positive for eosinophils (EPX) and the IL-33 receptor ST2 7 days after rIL-33 treatment. Inset, enlarged at bottom: left, red = EPX; middle, green = ST2; right = EPX, ST2 and nuclear marker dye Hoechst (blue) combined. Scale bars: Left top panel = 100μm, insets = 20μm. **b**, Absence of tumour regression after Myc-OFF in *KMG^null^* mice. Quantification of tumour burden in *KM* and *KMG* mice treated with tamoxifen (Myc ON), followed by cessation of treatment for two weeks (Myc OFF). *KM*: circular datapoints, *KMG GATA^null^*: square datapoints, *KMG GATA^het^*: triangular datapoints. Associated, right: representative H&E stains of whole lung sections for either condition / treatment. Scale bar: 2mm. n = 11 to 14 mice per condition. Statistical analysis was performed using the multiple Mann-Whitney tests. **c,** Induction of tumour cell death in *KMG^null^* mice. Quantification of immunohistochemical analysis for AT2 cell proliferation (Ki67/TTF1) and cell death (TUNEL) in *KM* and *KMG* mice treated with tamoxifen (Myc ON) or after Myc OFF. n = 6 mice per condition. Statistical analysis was performed using the multiple unpaired *t*-tests with Welch’s correction and based on the mouse averages. **b, c**: Data are from 2 ON/OFF, KM) or 3 (ON/OFF, KMG) independent biological replicate experiments.

## Discussion

Using our tamoxifen switch system to reversibly toggle MycER^T2^ activity in indolent KRas*^G12D^* + Myc driven lung LUAD we had previously shown that withdrawal or blockade of persistent Myc activity triggers rapid tumour regression^12^, consistent with previous reports in diverse switchable Myc mouse tumour models^44–48^. Myc-OFF-induced LUAD regression was characterised by rapid loss of Myc activity in LUAD cells (as evident from attenuation of expression of known Myc target genes), rapidly followed by a sharp fall in tumour cell proliferation and lesion burden, onset of tumour cell apoptosis, a dynamic regression choreography affecting all cell types within the tumour mass – epithelial, vascular, mesenchymal, inflammatory, and immune – and it concludes with the eventual normalisation of lung structure and vasculature. In our switchable mouse LUAD model, Myc is shut down only in the LUAD tumour cells. Hence, the rapid drop in Myc activity that triggers onset of regression must in some way be transduced out of the tumour cells and then propagated across the lung tumour milieu. To identify candidate pro-regression signals (ON and OFF) induced rapidly in LUAD tumour cells in response to Myc-OFF we used a combination of immune cytokine arrays and single-cell RNAseq. Both approaches converged on IL-33 as a compelling signal for relaying the Myc-OFF trigger that is immediately and transiently released from AT2-like LUAD cells upon Myc deactivation. We then established a causal role for IL-33 in driving LUAD regression by demonstrating both its necessity and sufficiency: antibody blockade of the (unique) IL33 receptor (ST2) profoundly inhibited Myc-OFF-induced LUAD regression. Conversely, systemic administration of a pulse of ectopic IL-33 to LUAD-bearing *KM* mice triggered LUAD regression, even if Myc activity was maintained throughout. Deactivation of Myc in lung tumours rapidly and transiently induces IL-33 release, likely as a result from chromatin disassociation and/or de novo synthesis^49^. This brief IL-33 elevation is sufficient for tumour regression and mirrors the dynamics of IL-33 as a damage sensor in response to lung injury^36,50^ during which IL-33 is transiently released from injured cells to restore homeostasis by engaging type 2 immunity cells^34,35,49,51^. These cells, in turn, promote the proliferation and differentiation of alveolar AT2 cells into gas-exchanging AT1 cells in addition to stimulating regeneration and repair of other—including structural—lung cell types^52–55^. Eosinophils express high levels of the IL-33 receptor ST2 and are obvious candidate effectors downstream of IL-33 in transduction of Myc-OFF-induced LUAD regression. Indeed, we observed that systemic administration of recombinant IL-33 in LUAD-bearing mice induced significant eosinophil influx in lungs. Using ΔdblGATA mice, which are deficient in development of eosinophils, Myc deactivation failed to drive regression of LUAD tumours, affirming eosinophils’ obligate role in IL-33-mediated transduction of the Myc-OFF regression programmes^35,49^. Like many other pleiotropic signalling molecules (such as TGFβ^56^), the roles that IL-33 plays in cancer remain both confusing and enigmatic, sometimes promoting cancer and at other times suppressing it, across tissues^57–64^. This dichotomy depends on tumour type, location, context, intensity and duration of the IL-33 signal and the target cell’s response. It also remains unclear whether IL-33’s role in tumour regression is coincident with its documented role in damage response or, instead, is just a convenient tissue-specific hook for evolution to use. Of note, we describe in an accompanying manuscript that acute downregulation of Myc in a mouse model of KRas*^G12D^*/MycER^T2^-driven pancreatic ductal adenocarcinoma (PDAC) also triggers a potent regression programme (Campos, 2026, manuscript in preparation). However, the sentinel PDAC cells release upon Myc-OFF is not IL-33 but GM-CSF, which is both necessary and sufficient for PDAC resolution and propagated thereafter by CD11c+ dendritic cells. The intriguing implication is that there may be substantial variety in the specific sentinel signals that mediate regression of tumour arising from different tissues. Our observations add further support the notion that targeting Myc in cancers—although difficult pharmacologically—retains several profound advantages over most other oncogenic targets. In both human and mouse studies, Myc blockade has surprisingly mild side effects, especially if applied metronomically. Mouse studies also indicate that partial blockade of Myc can be sufficient to thwart Myc oncogenic functions without appreciable harm to quiescent or continuously proliferating normal tissues^11,25,65^. Moreover, Myc is the unique, non-redundant conduit that sits at the bottom of the mitogenic signalling pathway and through which all upstream oncogenic signals (from kinases and G proteins) must flow. The guess is that cancer cells will have a much harder job rewiring their way past a Myc blockade. Our studies also offer a very different explanation for the long-standing mystery of so-called “oncogene addiction”: a foundational concept in targeted therapy that relies on acquired reliance for survival of cancer cells on a single dominant oncogene or signalling pathway, not just for continued growth but also survival. This includes Myc, even though it has no obvious role to play in maintaining cell survival. Many explanations for this phenomenon have been proffered, including asymmetry of attenuation of growth and apoptotic outputs in signalling^66,67^, constitutively high levels of intracellular stress signalling^68,69^ (oncogenic shock) or genetic instability^70^ in transformed tissues, terminal differentiation^45,71^, synthetic lethality and acquired interdependency between multiple oncogenic lesions^72^. Our data support a different interpretation: it is widely assumed that cancers are hacked and aberrantly persistent forms of the regenerative programme that repair wounds and injuries damaged tissues^1,73^. But wound repair has two major components—an initial regenerative phase, succeeded by resolution, a discrete morphogenic program that reorganizes the inchoate tissue and reasserts homeostatic architecture, cellularity, and function. Emerging evidence indicates that, in wound healing, the switch from regeneration to resolution is triggered by a decrease in mitogenic signalling (i.e. Myc). By analogy, in tumours blocking Myc, or oncogenic signals upstream of Myc, forcibly shuts down proliferation and forces engagement of resolution, the end result of which is orderly dismantling of the tumour and its regression.

## Material and Methods

### Mouse experiments and *in vivo* procedures

All treatments and procedures of mice were conducted in accordance with protocols approved by the Home Office UK guidelines under project licenses to G.I.E. (70/7586, PP2645677 and PP9124501) at the University of Cambridge and The Francis Crick institute. Mice were maintained on regular diet in a pathogen-free facility on a 7AM – 7PM 12hr light/dark cycle, IVC conditions 23°C and 61% humidity, with continuous access to food, water, and cage enrichment. *Kras^tm4Tyj^* (*Lsl-Kras^G12D^)* and *Gt(ROSA)26Sor^tm^*^1^(MYC/ERT2)*^Gev^ (LSL-Rosa26^MIE/MIE^)* mice have been described previously^74,75^. For all experiments the mice were kept with a heterozygous *Lsl-Kras^G12D^* allele and homozygous *LSL-Rosa26^MIE/MIE^*alleles. *Gata1^tm6Sho^ (*Δ*dblGATA)* mice^42^ were generously donated by Dr. Timothy Halim (CRUK Cambridge, UK) and are available at the Jackson Laboratories (RRID: IMSR_JAX:033551). They were crossed into the *Lsl-Kras^G12D^;Lsl-Rosa26^MIE/MIE^* background. Heterozygous and homozygous Δ*dblGATA* litters were confirmed by genotyping *Lsl-Kras^G12D^;Lsl-Rosa26^MIE/MIE^;*Δ*dblGATA* mice and used in experiments to assess the necessity of eosinophils in Myc OFF associated tumour regression. *B6(129S4)-Il33^tm^*^1^*^.1Bryc^/J* (*Il33^fl/fl^-eGFP*) mice^76^ were purchased from Jackson Laboratories (RRID: IMSR_JAX:030619) and crossed into the *Lsl-Kras^G12D^;Lsl-Rosa26^MIE/MIE^* background. *Il33^fl/fl^-eGFP* heterozygous and homozygous litters were confirmed by genotyping *Lsl-Kras^G12D^;Lsl-Rosa26^MIE/MIE^;Il33^fl/fl^-eGFP* mice and used in experiments to assess the necessity of AT2 tumour cell-derived IL-33 in Myc OFF associated tumour regression. For all experiments age and sex-matched littermate mice were divided equally over the treatment groups and *het/hom* genotypes. Mouse weight, health, and body condition was daily monitored during tamoxifen, blocking antibody or recombinant protein treatment. For activation of MycER^T2^, tamoxifen (Sigma; TS648) dissolved in peanut oil (Sigma; P2144) was administered daily by intraperitoneal (IP) injection for a maximum of 2 weeks at a dose of 1mg/25g body mass, or by tamoxifen diet (Harlan Laboratories UK, TAM400 diet) as previously described^77^ for a maximum of 4 weeks. Deactivation of MycER^T2^ was achieved by either cessation of tamoxifen IP injections or moving the mice from a tamoxifen diet back to a regular diet. To deliver adenovirus-Cre recombinase (AdV-Cre), 8 to 12 weeks old mice were anesthetised with isoflurane (Covetrus, Isofane 250 ml UK, 2800025; 100% w/w inhalation vapour with 0.2% oxygen) and 5×10^7^ plaque-forming units of AdenoCre (Ad5CMVCre, Viral Vector Core, University of Iowa) were administered as described previously^78^. For antibody blocking experiments during Myc-OFF driven tumour regression, mice were injected intraperitoneally (IP) with 150ug/IP of anti-ST2/IL-33R (monoclonal rat IgG2b clone 245707, MAB10041, R&D Systems) or isotype control (rat IgG2b, MAB0061, R&D Systems) every two days, starting two days before MycER^T2^ de-activation by cessation of tamoxifen injection. For systemic treatment of recombinant IL-33, after 2 weeks of tamoxifen food treatment (Myc ON) mice were IP injected for 2 days in a row (24 hours apart) with 1µg of recombinant mouse IL-33 (580504, BioLegend) or PBS control and then kept on tamoxifen food for another 3 or 7 days or 2 weeks (see Fig. 4, Extended Data Fig. 5) before euthanisation and lung collection. To stain tissues for hypoxia, 60 mg/kg hypoxyprobe-1 (1-([2-hydroxy-3-piperdinyl] propyl)-2-nitroimidazole hydrochloride) (Hypoxyprobe.com; HP1-100 kit) was administered through IP injection in saline 15 min prior to euthanasia. Protein adducts of reductively activated pimonidazole were identified through immunohistochemistry in paraffin-embedded fixed tissues with a monoclonal antibody against hypoxyprobe-1 (1:100). Lectin dye Rhodamine- *Ricinus communis* agglutinin I (RL-1082) was obtained from Vector Labs. Three minutes before sacrifice 100μl of a Rhodamine solution was administered by IV injection. Lungs were then harvested and fluorescence imaged on de-paraffinized tissue sections counterstained with Hoechst dye.

### Immunohistochemistry and immunofluorescence, multiplex RNAscope

Mice were euthanised and cardiac perfused with PBS. Lungs were removed, fixed overnight in neutral-buffered formalin (Sigma-Aldrich, 501320), and processed for paraffin embedding, Tissue sections (3 or 4 µm) were stained with haematoxylin and eosin (H&E) using standard reagents and protocols. For immunohistochemical analysis, sections were de-paraffinised, rehydrated, and boiled in a microwave for 10 minutes in citric acid-based antigen unmasking solution (H-3300, Vector labs) or universal heat induced antigen retrieval reagent (ab208572, Abcam), or treated for 15 minutes with 20μg/ml Proteinase K for antigen retrieval. Antibodies were incubated overnight at 4°C except for anti-CD3e from ThermoFisher, which was incubated for 20 minutes at room temperature. Antibodies used: polyclonal goat anti-MMR/CD206 (AF2535, Biotechne, 1:200); monoclonal rabbit anti-CD3e (SP7) (RM-9107-RQ, clone SP7, ThermoFisher, undiluted; ab16669, clone SP7, Abcam, 1:100); recombinant monoclonal rabbit anti-Ki67 (MA5-14520, Clone SP6, ThermoFisher 1:200); monoclonal rat anti-Ki67 (14-5689-82, Clone SolA15, ThermoFisher, 1:100); polyclonal rabbit anti-CD31 (ab28364, Abcam, 1:75); monoclonal rabbit anti-TTF1 (ab76013, clone EP1584Y, Abcam, 1:200); monoclonal rat anti-CD45R/B220 (MA1-70098, clone RA3-6B2, ThermoFisher, 1:100); monoclonal rat anti-CD335/NKp46 (137601, clone 29A1.4, BioLegend, 1:100); monoclonal rat anti-Ly6B.2 (NBP2-13077, clone 7/4, Novus Biologicals, 1:200); polyclonal goat anti-PDGFRa (AF1062, R&D Systems, 1:100); monoclonal rabbit anti-VWF (ab287962, clone EPR25069-131, Abcam, 1:100); monoclonal rabbit anti-IL-33 (ab187060, clone EPR17831, Abcam, 1:1000), monoclonal rat anti-IL-33 (MAB3626, clone 396118, R&D Systems, 1:100); monoclonal rat anti-ST2/IL-33R (MAB10041, clone 245707, R&D Systems, 1:100); monoclonal mouse anti-EPX kindly made available by E. Jacobson (MM25-82.2.1, Mayo Clinic Arizona, 1:5000). HRP-conjugated secondary antibodies (Vectastain Elite ABC Kits: PK-6200; Universal, PK-6101; Rabbit, PK-6104; Rat, PK-6105; Goat) were applied for 30 min and visualized with DAB (Vector Laboratories; SK-4100), or secondary Alexa Fluor 488 or -455 dye-conjugated antibodies (Life Technologies, see Reporting Summary for details) applied for 30 minutes at room temperature. DAB-stained slides were mounted with DPX mountant (Sigma, 06522) and fluorescence antibody-labelled slides were mounted in fluorescent mounting medium (Prolong Gold, Invitrogen, P36934) post-treatment with 0.5μg/ml Hoechst counterstain (Sigma-Aldrich, B2883). TUNEL analysis was performed using the Apoptag Fluorescein in situ Apoptosis Detection Kit (Millipore; S7110) according to manufacturer’s instructions. Briefly, tissue sections were pre-treated in Proteinase K (20μg/ml) for 15 min at room temperature, washed in deionized water twice for 2 min each, and returned to PBS. Sections were then covered with equilibration buffer for a minimum of 2 min followed by incubation at 37°C for 1 hr with a 1:5 dilution of TdT enzyme in reaction buffer. IHC and IF images for quantification were collected with a Zeiss Axio Imager.M2 microscope and Axiovision Rel 4.8 software (University of Cambridge), or with Zen3.2 (blue edition) software (Francis Crick Institute). RNAscope IF images and whole lung lobe imaging (see Extended Data Fig. 4a) were collected with a Vectra Polaris (Akoya Biosciences) imaging system and PotoImager HT software or a Zeiss Axio Scan.Z1 slide scanner with Zen3.7 (blue edition) software, respectively (Francis Crick Institute) and further processed withQuPath-0.5.1-x64.

### Sequencing

For bulk RNA sequencing, total RNA was isolated from *KM* lungs snap-frozen in liquid nitrogen using a Pure-Link RNA Mini Kit (12183018A, Thermo Fisher Scientific) and PureLink DNAse set (12185010, Thermo Fisher Scientific). The quality, quantity, and integrity of RNA were assessed by NanoDrop1000 spectrophotometer and 2100 Bioanalyzer (Agilent Technologies). RNA libraries were generated using TruSeq Stranded mRNA Library Prep Kit following the manufacturer’s instructions (20020594, Illumina, CA). RNA Libraries were run on the Ilumina NextSeq 500 using the 75-cycle high-output kit (single-end sequencing). The quality of the sequencing data was analysed by the bioinformatics team at the Cambridge Genomic Services (CGS) at the University of Cambridge (https://www.cgs.path.cam.ac.uk) using FastQC v0.11.4. In summary, reads were trimmed using Trim-Galore v0.5.0 and those less than 20 bases long were discarded. Reads were mapped using STAR v2.7.1. The Ensembl Mus Musculus GRCm38 (release 99) reference genome was used with annotated transcripts from the Ensembl Mus musculus GRC38.99.gtf file. The number of reads mapping to genomic features was calculated using HTSeq v0.6.1. Differential Gene Expression Analysis using the counted reads employed the R package edgeR v3.26.5 and the paired design model as suggested in the edgeR user’s guide. A GC and gene length correction using the CQN package (v 1.30.0) to remove systematic GC content and gene length bias via a smoothing function was applied. RNA-seq data have been deposited in the Gene Expression Omnibus database at NCBI (https://www.ncbi.nlm.nih.gov/geo) under accession number GSE312064. For single cell RNA sequencing, lungs were isolated from *KM* mice, placed into MACS tissue storage solution (130-100-008, Miltenyi) on ice and processed to single cell suspension on a gentleMACS Octo Dissociator with Heaters (130-096-427, Miltenyi) according to manufacturers’ instructions. In brief, after manually dissecting the lungs in ∼1mm^2^ pieces, they were further processed into a single cell suspension using the mouse tumour dissociation kit (130-096-730, Miltenyi) in gentleMACS C tubes (130-096-334, Miltenyi). After dissociation, the cell suspension was incubated with Red Blood Lysis Solution (130-094-183, Miltenyi) to remove red blood cells. Subsequently the cell suspension was filtered through 70µm MACS SmartStrainers (130-098-462) before being counted. 6000 cells per sample were offered for single cell sequencing. Library preparation of murine samples was performed according to instruction in the 10X Chromium single cell kit 3’ v3. The libraries were sequenced on a NovaSeq6000. Read processing was performed using the 10X Genomics workflow. Briefly, the Cell Ranger Single-Cell Software Suite (6.1.1) was used for demultiplexing, barcode assignment and UMI quantification (http://software.10xgenomics.com/single-cell/overview/welcome). The reads were aligned to the pre-built with the MYCER fusion gene sequence added. Raw single-cell sequencing data from individual samples were processed using the 10x CellRanger pipeline (10x Genomics) and analysed with the Seurat R-package (version 5.0.0)^79^ in R version 4.3.1. Cells were filtered to exclude cells with mitochondrial gene percentages higher than 25% and cells with less than 200 RNA features per cell. For sample integration, the R-package Harmony^80^ was used. In order to create the UMAP plots, the first 50 PCA dimensions were included, and cell-cycle regression was performed. The FindClusters function from the Seurat R-package was used to define clusters with the cluster parameter set to 0.5. The python package decoupler was used to assign cell type identities to clusters using the decoupler-py python package^81^ in conjunction with custom cell types and marker gene sets. Single-cell differential gene expression analyses were conducted with the glmGamPoi R-package^82^ (version 1.12.2). RNA-seq data have been deposited in the Gene Expression Omnibus database at NCBI (https://www.ncbi.nlm.nih.gov/geo) under accession number GSE311106.

### Immune Arrays

4 weeks post tamoxifen diet treatment of *KM* mouse (Myc ON), or 12 hours or 1, 2, 3, or 7 days after moving back to regular diet (Myc OFF), whole (tumour-laden) lungs were isolated and snap-frozen in liquid nitrogen. After crushing the frozen lungs into a powder using a mortar and pestle cooled with liquid nitrogen, the protein samples were isolated and incubated on a Proteome Profiler Mouse XL Cytokine Array (ARY028; Biotechne/R&D Systems) according to manufacturer’s instructions. For IF analysis and quantification array signals were visualised with IRDye 800CW Streptavidin secondary antibody (LI-COR, 926-32230). Array signals were captured and analysed using a LI-COR Odyssey CLx and Image Studio Software v5.0, respectively. Protein expression was calculated as relative to negative and positive assay controls included on the arrays.

### Quantification and statistical analysis of tumour burden, cytokine array, IL-33 protein expression, and immunohistochemistry

For quantifications of tumour burden, H&E sections were scanned with an Aperio AT2 microscope (Leica Biosystems) at 20X magnification (resolution 0.5 microns per pixel) and analysed with Image J. Data points on the dot plot graphs that portray tumour burden are represented as box-and whisker plots with range, which show upper extreme, upper quartile, median, lower quartile, lower extreme. Statistical significance was assessed by the unpaired non-parametric Multiple Mann-Whitney U test, with the median values calculated for each group. For the protein expression on the cytokine arrays (see Extended Data Figure 1b) the median values of 110 molecules of 8 or 9 whole (tumour-laden) lungs per timepoint/condition are displayed in a heatmap; CX3CL1 was excluded from the analysis because of unexplained gross variation in quantity dependent on experimental replicate, not sample identity. IHC staining was quantified by randomly selecting a minimum of 5 tumours per mouse and then counting the positively identified cells per tumour field of view (FoV). Data points on the scatter dot graphs that portray quantification per FoV represent one tumour (small data points) and averages per mouse (large data points), represented as box-and whisker plots with range, which show upper extreme, upper quartile, median, lower quartile, lower extreme; statistical significance was determined on the mouse means by unpaired Student’s t-test with Welch’s correction. For comparison, quantification of histological markers was only performed on tumour sections immunostained at the same time. For endothelial cell marker CD31 immunohistochemistry and the Ricinus communis rhodamine lectin dye the staining intensity was calculated as percentage CD31-positive or lectin-positive area per FoV using ImageJ. For proliferation marker Ki67 the immunofluorescence staining was quantified as percentage of Ki67-TTF1 double-positive cells using ImageJ. For quantification of NK cell marker NKp46 a complete full-face section of the lung of a mouse was analysed. Scoring (none, 10 cells, or > 10 cells percentage) was based on the amount of juxta-tumoral NKp46+ cells visible. Only tumours directly lining and associated with clearly distinguishable vasculature were taken into consideration; bar graphs are represented as mean with standard deviation; p-values are derived from mouse-average comparisons between groups and determined via two-way ANOVA with two degrees of freedom. All statistical significance was determined using Prism GraphPad software versions 9 and 10 (GraphPad). All graphs display p-values to four decimal places. The representative images represent at least five independent replicates.

### Sample sizes, biological replicates, blinding

No statistical methods were used to predetermine sample size. Mice were treated with blocking antibody for ST2 or with rIL-33 until tissue collection in a single-blind fashion. The investigators were not blinded during outcome assessment. Data in dot plot graphs is presented as the median and range (+/- upper and lower quartiles). Pilot experiments and previously published results were used to estimate the sample size. Mice euthanised before the end date of an experiment were excluded. Figure 1, Extended Data Figure 2: Cytokine arrays; median values are from three replicate experiments. n = 8 or 9 total per timepoint. Bulk RNA sequencing: n = 4 or 5 per timepoint. Singe cell RNA sequencing: Myc ON n = 8, from 4 replicate experiments, Myc OFF 2hrs n = 6, Myc OFF 4hrs n = 6, Myc OFF 8hrs n = 4, Myc OFF 12hrs n = 4, Myc OFF 24hrs n = 4. All Myc OFF samples are from two replicate experiments. Figure 2, Extended Data Figure 3: 1-week Myc OFF and ST2 blocking: n = 8 or 9 samples total per timepoint/condition, from 3 replicate experiments. 3 or 7 days Myc OFF and ST2 blocking; n = 8 or 9 samples total per timepoint/condition, from 2 (3d OFF) or 3 (7d OFF) replicate experiments. Figure 3, Extended Data Figure 4: Tumour burden Myc ON then OFF in *KMI* mice, *IL-33^fl/fl^-eGFP* heterozygous (*fl/+*) or homozygous (*fl/fl*) IL-33 floxed: Myc ON *fl/+* n = 2, *fl/fl* n = 7; Myc OFF *fl/+* n = 7, *fl/fl* n = 11, from 2 replicate experiments. Histochemistry Myc ON then OFF in *fl/fl* or *fl/+* mice for 3 or 7 days; n = 5 to 8 samples per condition/timepoint, from 2 replicate experiments, with 2 *fl/fl* samples and 4 *fl/+* samples for the Myc ON condition. Assessment of tumour IL-33 positivity in *KMI* mice: *fl/+* N = 9 mice, 2 Myc ON, 7 Myc OFF – total tumours analysed n = 445; *fl/fl* N = 12 mice, 5 Myc ON, 7 Myc OFF – total tumours analysed n = 751. Figure 4, Extended Data Figure 5: Tumour burden Myc ON 4 weeks +/- rIL-33; n = 7 to 10 samples per condition/timepoint, from 3 replicate experiments. Tumour burden Myc ON +/- rIL-33 3 or 7 days; n = 7 to 10 samples per condition/timepoint, from 3 replicate experiments. Histochemistry Myc ON +/- rIL-33 3 or 7 days; n = 5 or 6 samples per condition/timepoint, from 3 replicate experiments. Recombinant IL-33 in *KM* (AdVCre and Tamoxifen treated (3 replicate experiments) versus AdVCRE and tamoxifen untreated (one replicate) mice, representative of 6 mice per condition. Figure 5, Extended Data Figure 6 : Tumour burden Myc ON then OFF in *KM* or *KMG* mice, Δ*dblGATA* heterozygous (*+/-*) or homozygous (*-/-*) null: Myc ON *KM* n = 6, KMG^null^ n = 8 – total n = 14; Myc OFF *KM* n = 8, *KMG^het^* n = 3, *KMG^null^* n = 12 – total 23 mice, from 2 (ON/OFF *KM*, or 3 (ON/OFF *KMG*) replicate experiments. Histochemistry Myc ON then OFF in *KM* or *KMG*; n = 6 samples per timepoint, equally divided in Myc ON *KM* n = 3 and KMG^null^ n = 3, or Myc OFF *KM* n = 3, *KMG^het^* n = 3, while *KMG^null^* n = 6, from 2 (ON/OFF *KM*), or 3 (ON/OFF *KMG*) replicate experiments.

### Data availability

Raw and processed bulk RNA-seq and single cell–seq data generated in this study were deposited in the NCBI Gene Expression Omnibus (accession no. GSE311106 and GSE312064, respectively). All other relevant data supporting the key findings of this study are available within the Article and its Extended Data Figures or from the corresponding author upon reasonable request.

## Acknowledgments

We thank the members of the Evan laboratory for discussion and the BRU staff at the Cambridge Cancer Research Institute (CRUK CI) and BRF staff at the Francis Crick Institute for expert animal care. We also thank Abigail Edwards and staff at the Pre-clinical Genome Editing core at the CRUK CI and Lucy Meader at the Histopathology Core at the Francis Crick Institute for their expert support. This study was supported by programme grants to GIE (Cancer Research UK C4750/A12077).

## Author contributions

RMK and GIE conceived the project and designed experiments. RMK supervised and performed all experiments. TC and KU-J assisted with some experiments. SB performed bioinformatic analysis. AP performed all mouse experimental procedures. RMK and GIE wrote the manuscript. GIE supervised the study. All authors discussed results and revised the manuscript.

## Additional information

Correspondence and request for material can be sent to R.M. Kortlever and G.I. Evan.

---

**Extended Data Figure 1. Schematic representations of animal experiments a**, Myc ON 4 weeks then OFF 2 or 4 hours – Bulk RNA sequencing

**b**, Myc ON 4 weeks then OFF 2, 4, 8, 12 or 24 hours – Single cell RNA sequencing

**c**, Myc ON 4 weeks then OFF 12 hours or 1, 2, 3, 7 days – Protein Cytokine Array **d**, Myc ON 2 weeks then OFF 3 or 7 days plus anti-ST2 antibody or IgG control **e**, Myc ON 2 weeks then OFF 3 or 7 days in *Il33fl/+* or *Il33fl/fl* mice

**f**, Myc ON 2 weeks then treatment with recombinant IL-33 (rIL33) or PBS control followed by a further 2 weeks Myc ON

**g**, Myc ON 2 weeks then treatment with recombinant IL-33 (rIL33) or PBS control followed by a further 3 or 7 days Myc ON

**h**, Myc ON 2 weeks then treatment with recombinant IL-33 (rIL33) followed by a further 7 days Myc ON versus age-controlled mice without tumour activation and recombinant IL-33 treatment followed by a further 7 days of no Myc ON

**i**, Myc ON 2 weeks then OFF 2 weeks in *ΔdblGATA* (eosinophil KO) mice For all experiments: blue triangles indicate lung collection timepoints.

**Extended Data Figure 2. Acute Myc loss immediately reverses Myc-regulated gene expression**

**a**, Myc-OFF very rapidly modifies Myc target gene expression. Principal Component Analysis (PCA) on Bulk RNA-sequence data for Hallmark Myc targets (M5926) of the Myc ON and 2 or 4 hours Myc OFF timepoints, and KRas*^G12D^*-only. Data are from n = 4 or 5 independent mouse lungs per timepoint/condition.

**b**, Myc-OFF immediately reverses Myc target gene expression. Bulk sequencing analysis of tumour laden lungs 2 and 4 hours after Myc OFF, compared to Myc ON. Shown is a heatmap of a curated MycER^T2^ target list specific for *KM* mouse and ranked according to most change compared to KRas*^G12D^-*only (not shown) when Myc ON. n = 4 or 5 mice per timepoint/condition. Gene list is shown in Extended Data Table 1.

**c**, Myc-OFF does not immediately change non-Myc target gene expression. Analysis as in (**b**), with a curated housekeeping-associated gene list, where the ranked Myc ON Log_2_ Fold Change of expression was non-significant, between -0.1 to 0.1 times of KRas*^G12D^*-only, and ranked as induced (tangerine colour) or down (grape colour) in the first column of the heatmap. For comparison with (**b**) the equivalent Log_2_ Fold Change range is depicted. For the two gene collections (**b** and **c**) genes were used with measurable expression for all four conditions. n = 4 or 5 mice per condition. Gene list is shown in Extended Data Table 1.

**d**, Rapid reversion of Hallmark gene sets expression after Myc-OFF. Normalised enrichment score (NES) of the top-scoring Hallmark gene sets in the molecular signatures database (MSigDB) after gene set enrichment analysis. Left graph: positively or negatively related with switching Myc ON 4 weeks versus KRas*^G12D^*-only. Right graph: positively or negatively related with switching Myc OFF for 4 hours versus Myc ON. Within the bars are mentioned the significant FDR *q*-values. NES was calculated by weighted Kolmogorov-Smirnov test and FDR *q*-value was determined by permutation-based testing with multiple Benjamini-Hochberg hypothesis correction. n = 4 or 5 mice per timepoint/condition.

**e**, Cell type marker expressions in Myc ON then OFF single cell experiment. Matrix plot depicting unique cluster/cell-type specific gene identifiers from experimental procedure depicted in Extended Data Figure 1b.

**f**, UMAP representation of spatiotemporal Myc ON then OFF experiment. Single-cell suspensions of 32 tumour-laden whole mouse lungs from 6 different timepoints before and after Myc OFF were isolated and analysed by RNA-sequencing (see Extended Data Figure 1b). UMAP representation of cells that were annotated and grouped by colour into 30 different clusters. See Extended Data Table 2 for full annotations. n = 32 mice total; 4-8 mice per timepoint, with 6 timepoints in total, from up to 4 independent biological replicate experiments. **g**, Myc-OFF instantly reverses Myc-target gene expression in MycER^T2^-positive tumour cells. Gene set enrichment analysis (GSEA) of Hallmark Myc-targets_V1 (M5926) in *MycER^T^*^2^ positive cells from single cell analysis after Myc OFF 2 hours (see Figure 1b, Extended Data Figure 1b, Extended Data Figure 2e, f). Normalised enrichment score (NES) was calculated by weighted Kolmogorov-Smirnov test and FDR *q*-value was determined by permutation-based testing with multiple Benjamini-Hochberg hypothesis correction. n = 8 (Myc ON) or 4 (2hrs Myc OFF) from 4 (Myc ON) or 2 (2hrs Myc OFF) biological replicate experiments.

**Extended Data Figure 3. Immediate and temporal IL33 expression after acute Myc loss a**, Immune signalling molecule expression analysis in the first week of Myc-OFF in LUAD. Heatmap representation of immune cytokine arrays of whole tumour-laden lungs at various timepoints after Myc OFF versus Myc ON. Protein expression of 110 immune-associated molecules was normalised for Myc ON and subsequently ranked in order of fold change at 12hrs Myc OFF. Up- and downregulated proteins are shown in tangerine and grape colour, respectively. Every column shows the median protein expression values for n = 8 or 9 mice. Data are from three independent biological replicate experiments.

**b**, IL33 expression in LUAD first 3 days after Myc-OFF. Quantification of IL-33 protein expression as analysed by immune cytokine array over serial timepoints after Myc OFF. n = 8 or 9 mice per timepoint/condition. Statistical analysis was performed using the multiple unpaired *t*-tests with Welch’s correction.

**c**, *Il33* expression in MycER^T2^-positive epithelial tumour cells. Violin plot representations of *Il33* and *MycER^T^*^2^ expression in the four main epithelial cell type clusters. n = 32 mice total; 4-8 mice per timepoint, with 6 timepoints in total, from up to 4 independent biological replicate experiments.

**Extended Data Figure 4. IL-33 Signalling regulates the immune and neoangiogenic microenvironment**

**a**, Blocking IL33 receptor ST2 after Myc-OFF prevents vascular normalisation and hypoxia. Quantification of *Ricinus communis* lectin dye staining (top graph) and quantification of immunohistochemical Hypoxyprobe staining (bottom graph); analysis of vascular changes of lung tumours in *KM* mice treated with tamoxifen (Myc ON) followed by Myc OFF for 3 days while simultaneously treated with anti-ST2 or IgG control. Panels to their right: representative immunohistochemistry. n = 4 mice per condition/timepoint. Scale bar 50μm (top graph) or 100μm (bottom graph). Data are from one experiment.

**b**, Blocking IL33 receptor ST2 after Myc-OFF prevents some but not all immune cell changes. Quantification of immunohistochemical analysis for B cell (B220), eosinophil (EPX), and neutrophil (Ly6B.2) presence in lung tumours in *KM* mice treated with tamoxifen (Myc ON) followed by Myc OFF for 3 or 7 days while simultaneously treated with anti-ST2 or IgG control antibody. FoV = field of view. Panels to their right: representative immunohistochemistry. n = 6 to 9 mice per condition or timepoint. Scale bars 100μm. Data are from 2 (3 days Myc OFF) or 3 (7 days Myc OFF) independent biological replicate experiments.

**Extended Data Figure 5. IL-33 from tumour AT2 cells is necessary for Myc OFF induced reversal of immune suppression**

**a**, Efficiency of conditional loss of IL33 in *KMI* mice. Qualitative representations of immunofluorescence staining of mice used in experiment depicted in Extended Data Fig. 1e and Fig. 3a. Left column shows protein IL-33 and the nucleus (Hoechst), right column AT2 cell marker TTF1 and proliferation marker Ki67. Shown is part of the major lung lobe, see top right H&E: IF outline = Immunofluorescence. Top and bottom rows: *KMI^fl/fl^* mice, middle row: *KMI^fl/+^*mice. Top row: Myc ON, bottom two row: Myc OFF. Insets are enlarged representations of white outlined squares in every panel. Scale bars: 500μm (large panels), or 100μm (insets).

**b**, Tumour immune profiling of *KMI^fl/+^* versus *KMI^fl/fl^* mice after Myc-OFF. Quantification of immunohistochemical analysis for Macrophages (CD206), T cells (CD3), or eosinophils (EPX) in lung tumours of *KMI^fl/+^* or *KMI^fl/fl^* mice treated with tamoxifen (Myc ON) followed by Myc OFF for 3 days. n = 5 or 6 mice per condition and timepoint. Statistical analysis was performed using the multiple unpaired *t*-tests with Welch’s correction and based on the mouse averages. **c**, Loss of tumour IL33 after Myc-OFF prevents NK cell accretion. Quantification of immunohistochemical analysis for juxta-tumoural NK cell (NKp46) presence in *KMI^fl/+^*or *KMI^fl/fl^* mice treated with tamoxifen (Myc ON) followed by Myc OFF for 3 days. n = 5 or 6 mice per condition and timepoint. Statistical analysis was performed using the two-way ANOVA with Tukey’s multiple comparisons test.

**b, c**: Data are from 2 independent biological replicate experiments.

**Extended Data Figure 6. Transient systemic IL-33 treatment induces instant tumour regression**

**a**, rIL33 imposes very rapid tumour resolution. Quantification of tumour burden in *KM* mice without treatment (KRas*^G12D^-*only), treated with tamoxifen (Myc ON) and PBS control or rIL-33 and collected 3 or 7 days after treatment while continued Myc ON. Statistical analysis was performed using the multiple Mann-Whitney tests. Data are from 3 independent biological replicate experiments.

**b**, rIL33 imposes eosinophil influx into lung regardless of tumour presence. H&E or immunohistochemistry of tumour and non-tumour lung tissue 7 days post rIL-33 treatment. Left, H&E staining: whole lung sections per treatment (top panels: *KM* tumours, bottom panels: *KM* no tumours), plus enlargements (two insets per condition) of one small part of one lung lobe (insets, bottom row) to highlight remnant lesions and/or eosinophil aggregation. n = 7 to 10 mice per condition. Scale bars 2mm (whole tissue section); 200μm (bottom left inset) or 50μm (bottom right inset). Right: representative immunohistochemistry (IHC) analysis of AT2 cell (TTF1) and eosinophil (EPX) presence in selected inset sections from the H&E staining shown at their left. Scale bar 100μm. Data are from 3 (AdenoCRE) or 1 (No AdenoCRE) independent biological replicate experiments.

**Extended Data Figure 7. Eosinophils are necessary for reversal of immune cell mobilisation after Myc OFF**

**a**, Eosinophils are required for tumour-local Myc-OFF driven immunity. Quantification of immunohistochemistry analysis for macrophages (CD206) and T cells (CD3) in lung tumours of *KM* mice treated with tamoxifen (Myc ON) followed by regular food (Myc OFF) for 2 weeks. FoV = field of view. n = 6 mice per condition. Statistical analysis was performed using the multiple unpaired *t*-tests with Welch’s correction and based on the mouse averages.

**b**, Absence of eosinophils after Myc-OFF prevents NK cell attraction. Quantification of immunohistochemistry analysis for juxta-tumoural NK cells (NKp46) of lung tumours in *KM* mice treated with tamoxifen (Myc ON) followed by regular food (Myc OFF) for 2 weeks. n = 6 mice per condition. Statistical analysis was performed using the two-way ANOVA with Tukey’s multiple comparisons test.

**a, b**: Data are from 2 ON/OFF, KM) or 3 (ON/OFF, KMG) independent biological replicate experiments.

