## Supplemental Figures for "Targeting Myc activates a tissue-specific tumour resolution programme"

### Extended Data Figure 1. Schematic representations of animal experiments

**a** Related to Extended Data Figure 1a-d

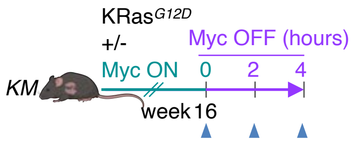


**b** Related to Figure 1b, Extended Data Figure 1e-g, 3c

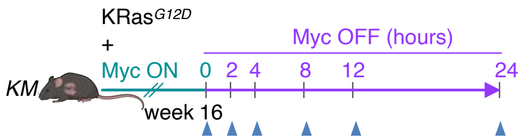


**c** Related to Figure 1a, Extended Data Figure 3a, b

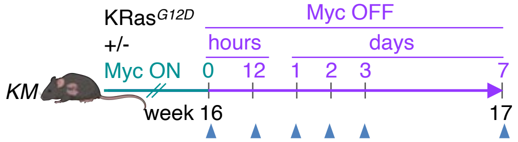


**d** Related to Figure 2a-d, Extended Data Figure 4a, b

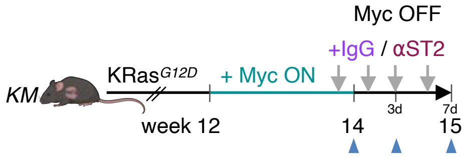


**e** Related to Figure 3a-d, Extended Data Figure 5a-c

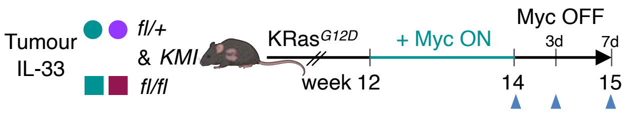


**f** Related to Figure 4a

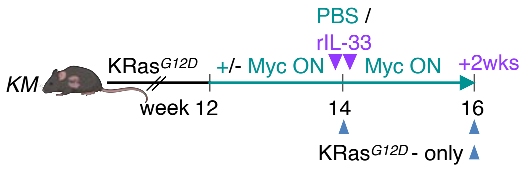


**g** Related to Figure 4b-e, 5a,,b, Extended Data Figure 6a

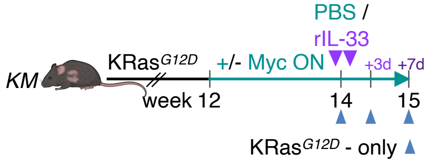


**h** Related to Extended Data Figure 6b

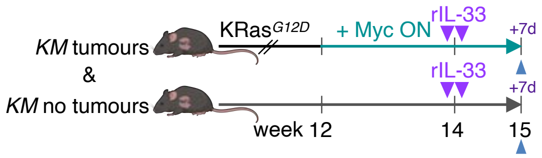


**i** Related to Figure 5b, c, Extended Data Figure 7a, b

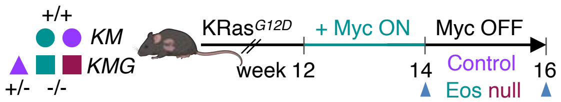

Extended Data Figure 2. Acute Myc loss immediately reverses Myc-regulated gene expression

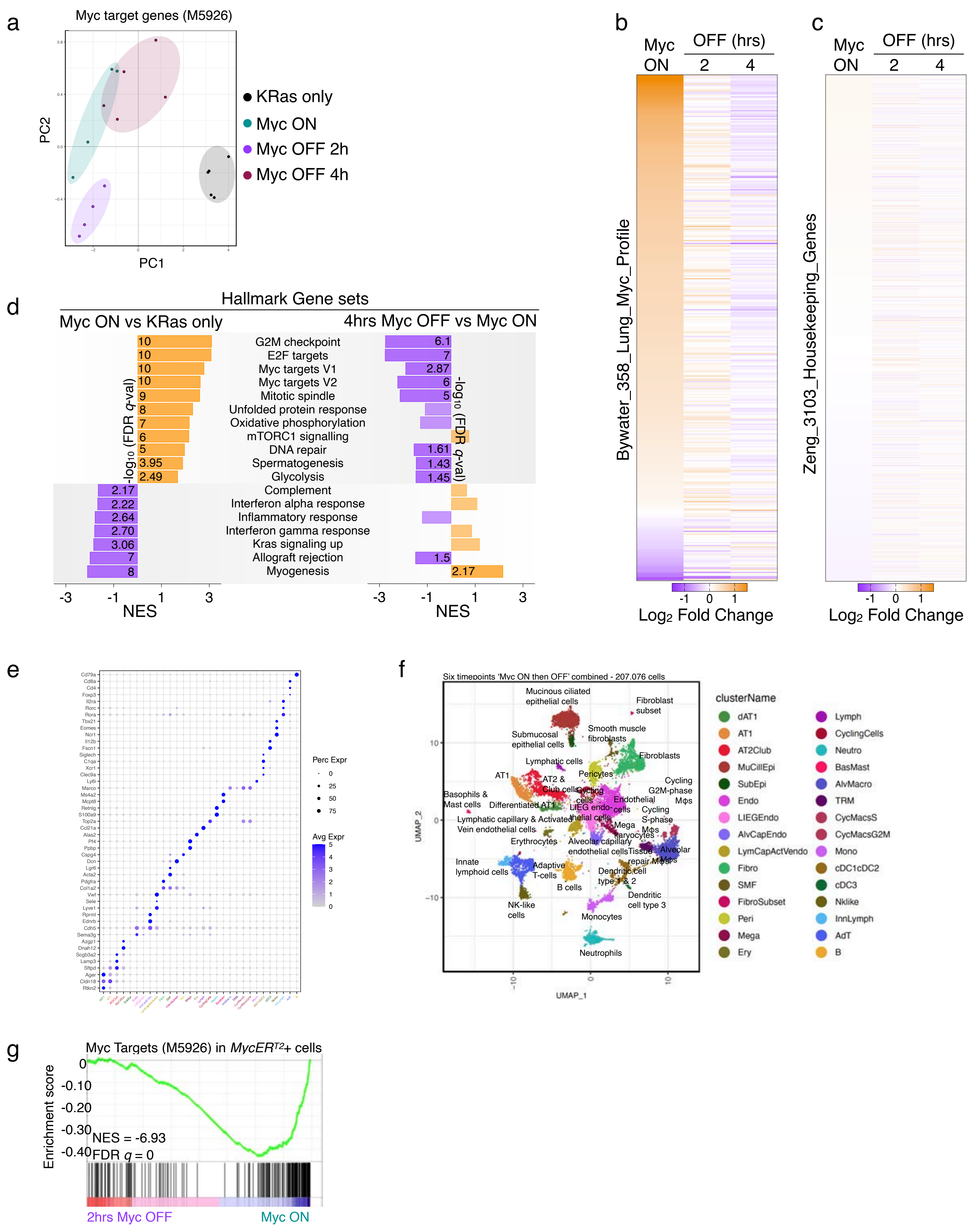

Extended Data Figure 3. Immediate and temporal IL33 expression after acute Myc loss

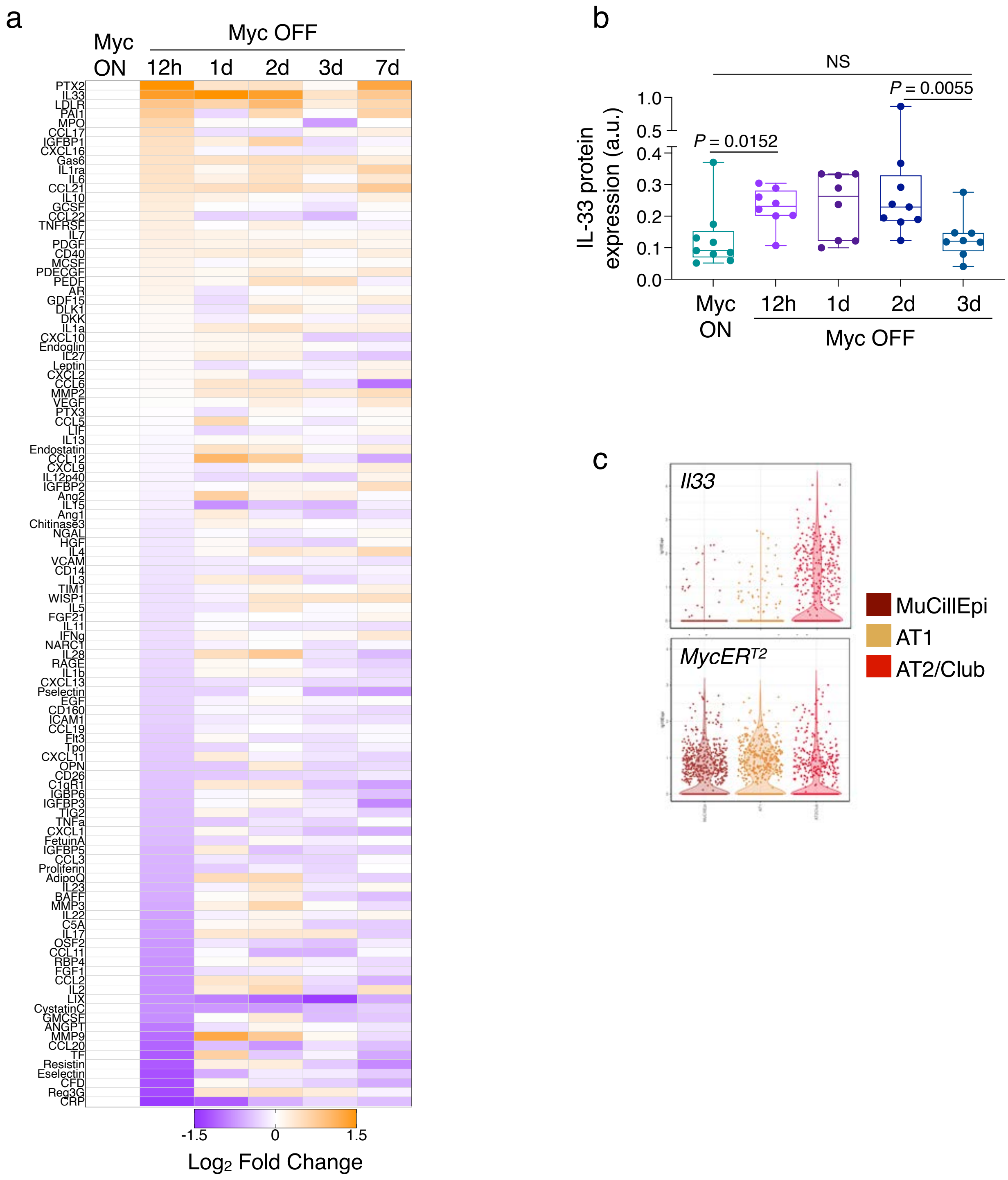

Extended Data Figure 4. IL-33 Signalling regulates the immune and neoangiogenic microenvironment

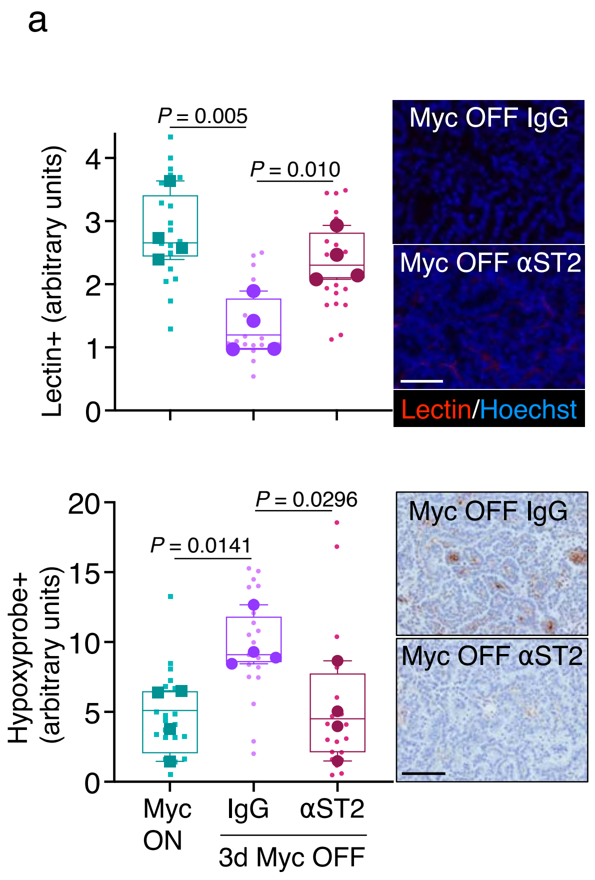


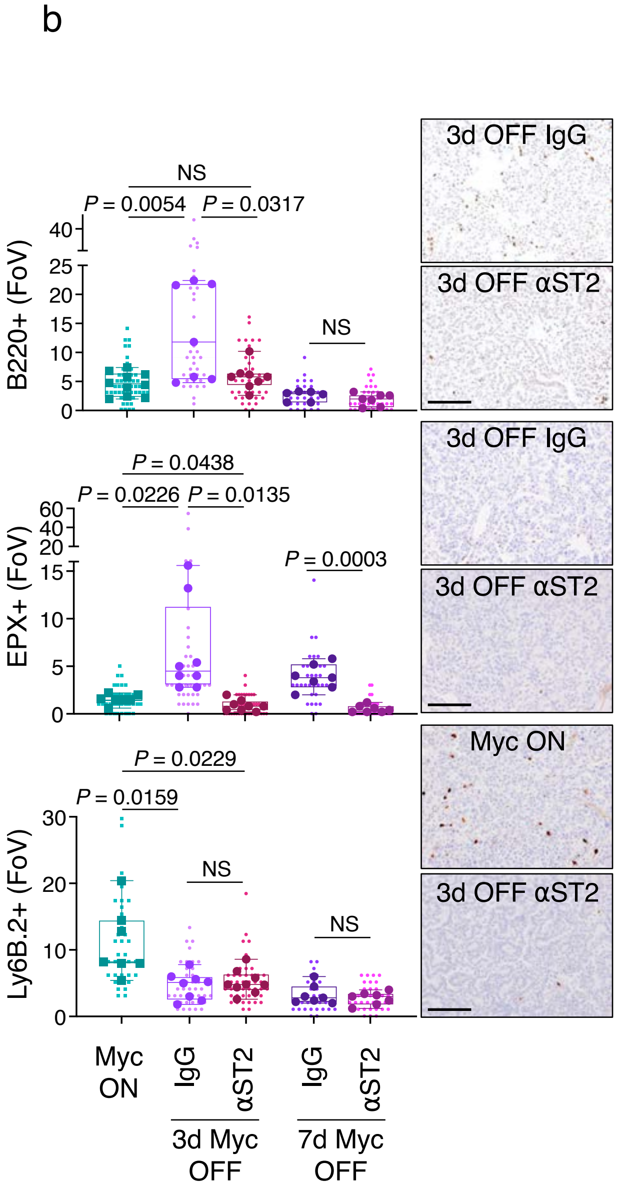

Extended Data Figure 5. IL-33 from tumour AT2 cells is necessary for Myc OFF induced reversal of immune suppression

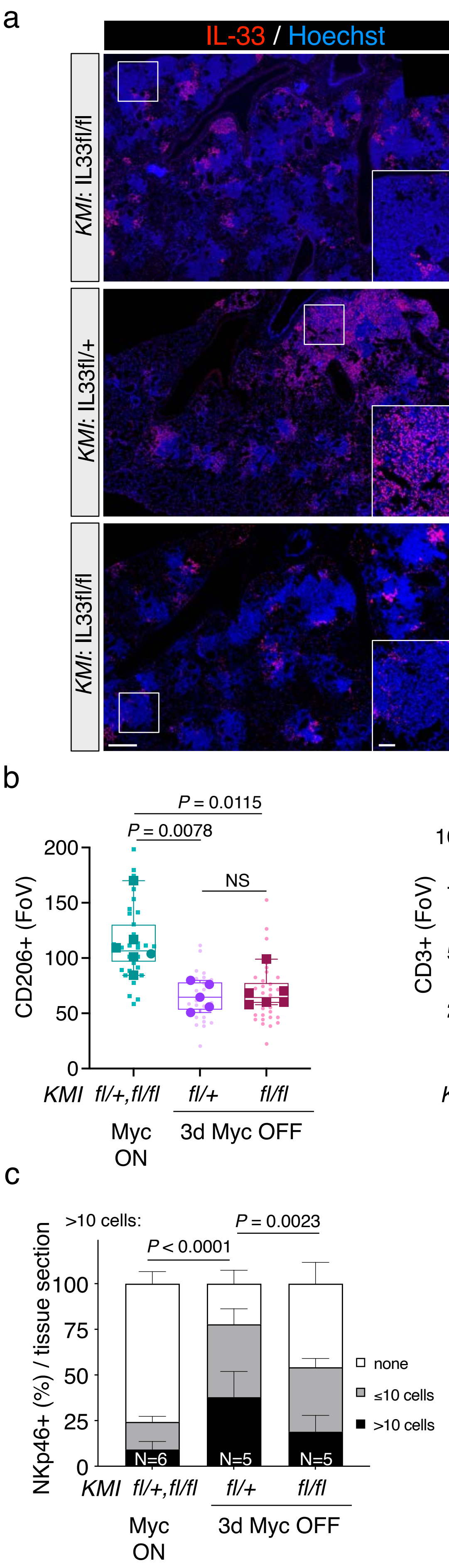

Extended Data Figure 6. Transient systemic IL-33 treatment induces instant tumour regression

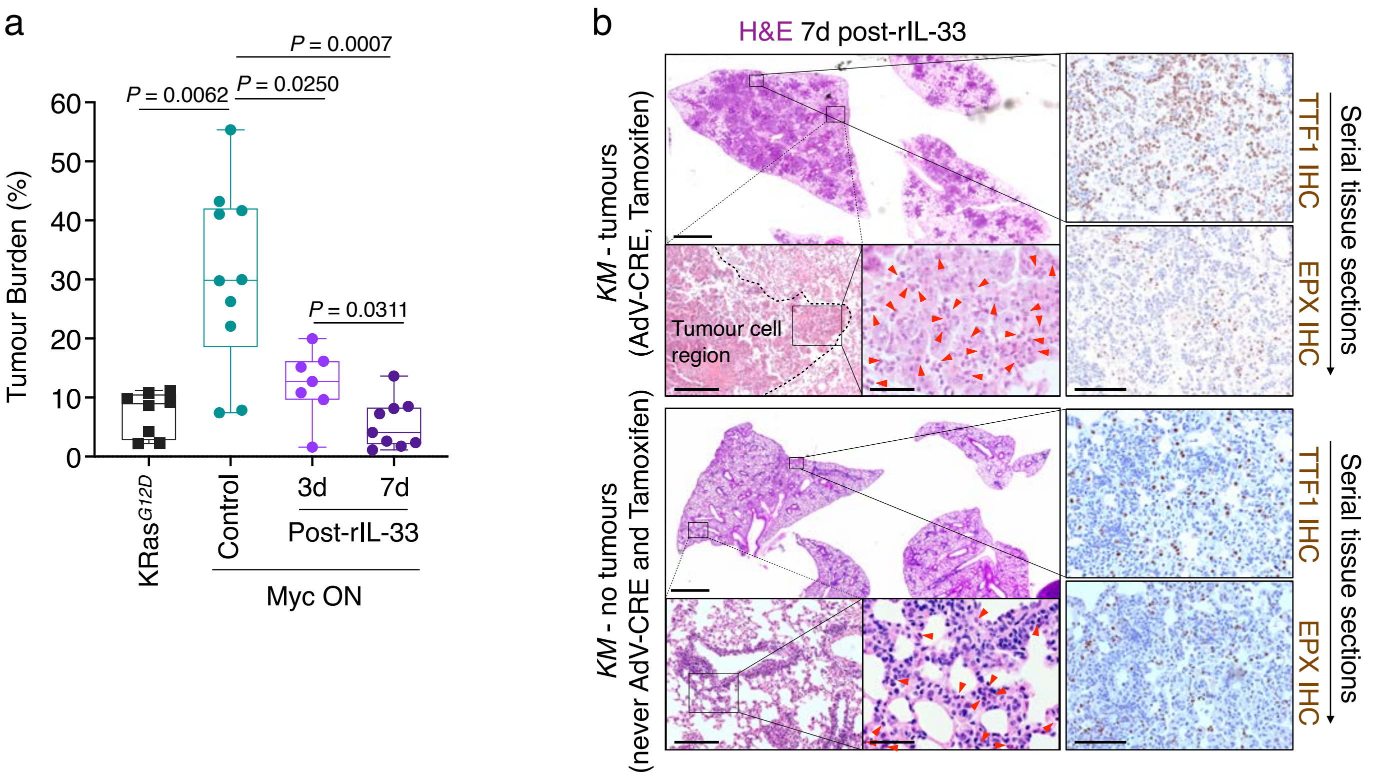

Extended Data Figure 7. Eosinophils are necessary for reversal of immune cell mobilisation after Myc OFF

a

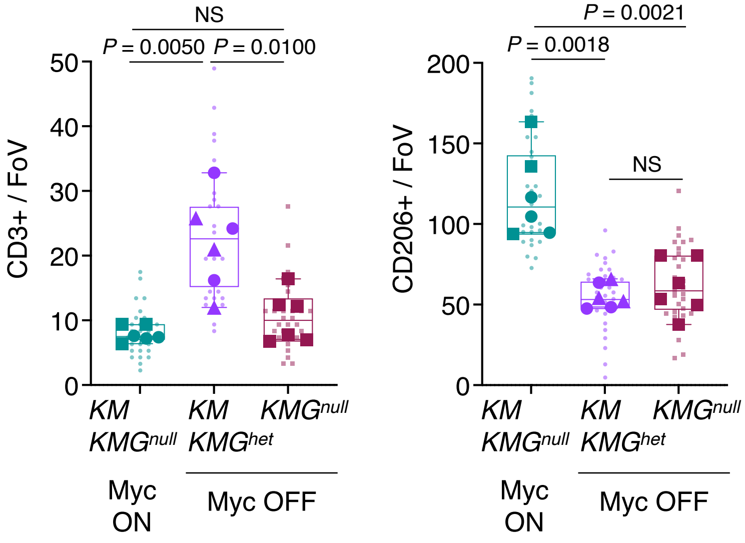


b

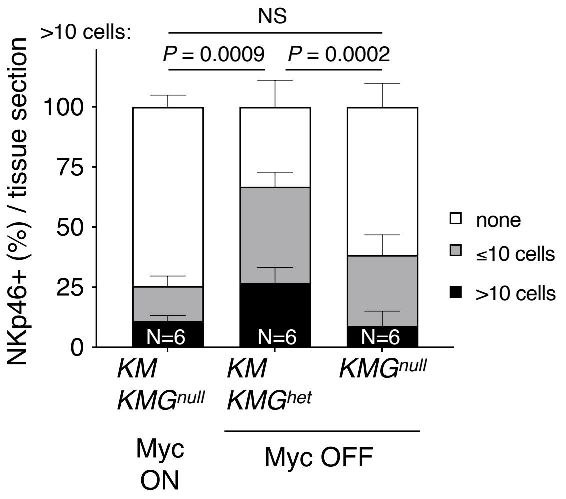
